# A phylogenetic and proteomic reconstruction of eukaryotic chromatin evolution

**DOI:** 10.1101/2021.11.30.470311

**Authors:** Xavier Grau-Bové, Cristina Navarrete, Cristina Chivas, Thomas Pribasnig, Meritxell Antó, Guifré Torruella, Luis Javier Galindo, Bernd Franz Lang, David Moreira, Purificación López-Garcia, Iñaki Ruiz-Trillo, Christa Schleper, Eduard Sabidó, Arnau Sebé-Pedrós

## Abstract

Histones and associated chromatin proteins have essential functions in eukaryotic genome organization and regulation. Despite this fundamental role in eukaryotic cell biology, we lack a phylogenetically-comprehensive understanding of chromatin evolution. Here, we combine comparative proteomics and genomics analysis of chromatin in eukaryotes and archaea. Proteomics uncovers the existence of histone post-translational modifications in Archaea. However, archaeal histone modifications are scarce, in contrast with the highly conserved and abundant marks we identify across eukaryotes. Phylogenetic analysis reveals that chromatin-associated catalytic functions (e.g., methyltransferases) have pre-eukaryotic origins, whereas histone mark readers and chaperones are eukaryotic innovations. We show that further chromatin evolution is characterized by expansion of readers, including capture by transposable elements and viruses. Overall, our study infers detailed evolutionary history of eukaryotic chromatin: from its archaeal roots, through the emergence of nucleosome-based regulation in the eukaryotic ancestor, to the diversification of chromatin regulators and their hijacking by genomic parasites.

## Introduction

The access to genetic information in eukaryotes is controlled by a manifold nucleoproteic interface called chromatin. This nucleosomal chromatin environment defines a repressive ground state for transcription and other DNA-templated processes in eukaryotic genomes^1,2^. Multiple components associated with chromatin underlie elaborate eukaryotic genome regulation, allowing the differential access to genetic information in time/space and the maintenance of the resulting regulatory states^3–6^. Moreover, chromatin-based regulation is essential in repressing parasitic genomic elements, like transposons and viruses^7–11^.

The main protein components of eukaryotic chromatin are histones. All eukaryotes have four major types of histones (H2A, H2B, H3 and H4), which are combined as an octamer to form the basic repetitive unit of the chromatin: the nucleosome. Canonical histones are among the most highly conserved proteins across eukaryotes^12^ and, in addition, unique histone variants (paralogs of one of the four major histone types) are found in many species, often associated with particular regulatory states^13–17^. Histone chemical modifications, including acetylations and methylations play a central role in genome regulation and transgenerational epigenetic inheritance^3,18–21^. These chemical moieties, known as histone post-translational modifications (hPTMs), are added and removed by specific enzymes (‘writers’, e.g., histone methyltransferases or acetylases; and ‘erasers’, e.g., histone demethylases and deacetylases). Some hPTMs (e.g., most acetylations) have a generic effect on nucleosome stability, while others are bound by specific proteins or protein complexes. These are often referred to as ‘readers’ and include proteins like HP1, which binds to H3K9me3, as well as a myriad of other proteins encoding Chromo, PHD, Tudor and Bromo structural domains, among others^22–24^. Finally, nucleosome remodellers (like SNF2 proteins) and histone chaperones are additional important players in chromatin regulation, by mediating chromatin opening, nucleosomal assembly, and histone variant interchanges^25–28^.

All eukaryotes studied to date possess histone-based chromatin organization, with the sole exception of dinoflagellates, which nonetheless encode for histone proteins in their genomes^29^. Beyond eukaryotes, histones have also been identified in Archaea, where they have been shown to form nucleosomal structures^30–33^. However, unlike eukaryotic histones, the few archaeal histones experimentally characterized so far (*i*) generally lack disordered *N*-terminal tails; (*ii*) do not have any known post-translational modifications^34^; and (*iii*) do not seem to impose a widespread, genome-wide repressive transcriptional ground state^33,35^. Thus, chromatin-based elaborate genome regulation is often considered a eukaryotic innovation^36,37^.

From a phylogenetic perspective, our understanding of chromatin components and processes derives from a very small set of organisms, essentially animal, fungal and plant model species plus a few parasitic unicellular eukaryotes. Additional efforts have sampled specific aspects of chromatin regulation, such as histone modifications or their genome-wide distribution, in non-model animal species^38,39^, fungi (*Neurospora crassa* and *Fusarium graminearum*)^40,41^, and five other eukaryotes: the unicellular holozoan *Capsaspora owczarzaki*^42^, the dinoflagellate *Hematodiunium* sp.^29^, the brown alga *Ectocarpus siliculosus*^43^, the amoebozoan *Dictyostelium discoideum*^44^, and the ciliate *Tetrahymena thermophila*^45,46^. However, these organisms represent a tiny fraction of eukaryotic diversity. Hence, we lack a systematic understanding of the evolution of eukaryotic chromatin modifications and components^47^.

In order to infer the origin and evolutionary diversification of eukaryotic chromatin, we performed a joint comparative analysis of histone proteomics data from 30 different eukaryotic and archaeal taxa, including new data for 23 species. In parallel, we analyzed the complement of chromatin-associated gene families in an additional 172 eukaryotic genomes and transcriptomes. This comprehensive taxon sampling includes representatives of all major eukaryotic lineages, as well as multiple free-living members of enigmatic early-branching eukaryotes (e.g., jakobids, malawimonads, *Meteora* sp. and ancyromonads, as well as Collodictyonida, Rigifilida and Mantamonadida (CRuMS); **Fig. 1a**). In addition, in order to trace the pre-eukaryotic origins of these chromatin gene families, we systematically searched for orthologs in archaeal, bacterial and viral genomes. Specifically, we reconstructed the evolutionary history of enzymes involved in chromatin modification and remodelling; as well as the conservation of the hPTMs effected by these enzymes. Our comparative genomics and proteomics suggest a concurrent and early origin of canonical histones, a core of quasi-universal hPTMs, and their corresponding enzymatic effectors. We also identify independent expansions in hPTM reader gene families across eukaryotes and document evidence of the capture of these reader domains by parasitic genomic elements. Overall, this work provides a phylogenetically-informed framework to classify and compare chromatin components across the eukaryotic tree of life, and to further investigate the evolution of hPTM-mediated genome regulation.

**Figure 1.**
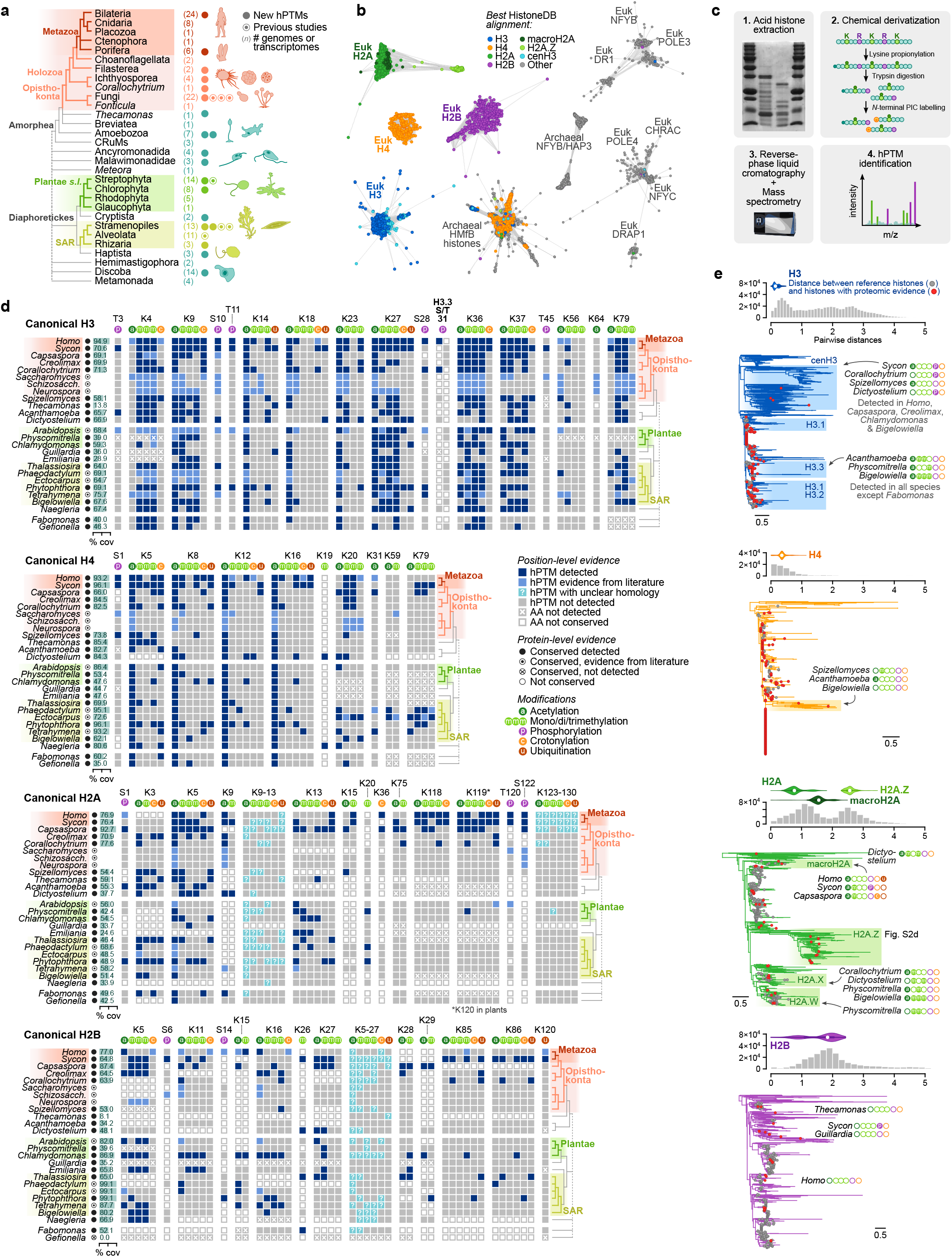
Diversity of post-translational modifications in eukaryotic canonical and variant histones. **a,** Eukaryotic taxon sampling used in this study. Colored dots indicate the number of species used in the comparative histone proteomics reconstruction, with solid dots indicating new species added in this analysis. Numbers in brackets indicate the number of genomes/transcriptomes used in the comparative genomics analyses. Dashed lines indicate uncertain phylogenetic relationships. Complete list of sampled species in Supplementary Table 1. **b,** Networks of pairwise protein similarity between histone protein domains in eukaryotes, archaea and viruses. Each node represents one histone domain, colored according to their best alignment in the HistoneDB database (see Methods). Edges represent local alignments (bitscore ≥ 20). **c,** Schematic representation of the hPTM proteomics strategy employed in this study. **d,** Conservation of hPTMs in eukaryotic histones. hPTM coordinates are reported according to the amino-acid position in human orthologs (if conserved). In H2A and H2B, question marks indicate the presence of hPTMs in stretches of lysine residues of uncertain homology. In species with previously reported hPTMs, we further indicate which variants were also identified in our reanalysis. Only positions with hPTMs conserved in more than one species are reported (full table and consensus alignments available in Supplementary Table 3). **e,** Maximum likelihood phylogenetic trees of the connected components in panel b, corresponding to eukaryotic histones (H3, H4, H2A, H2B). Canonical histones included in panel d and variant histones detected are highlighted in red. hPTMs detected in non-canonical histones are indicated. Bottom, distributions of pairwise phylogenetic distances between all proteins in each gene tree. Violin plots above each distribution represent the distribution of distances between reference histones present in the HistoneDB database and histones with proteomic evidence included in our study, for each of the main canonical (H3, H4, H2A, and H2B) and variant histones (H2A.Z and macroH2A).

## Results

### Comparative proteomics of eukaryotic histone post-translational modifications

We analyzed the phylogenetic distribution and evolutionary history of histone proteins. To this end, we surveyed the presence of histone-fold proteins across 172 eukaryotic and 4,226 archaeal taxa, using HMM searches (**Fig. 1a,b and Supplementary Table 1**). Histone proteins are found in all eukaryotic genomes. We clustered the identified 8,576 histone-encoding proteins using pairwise local alignments and then classified individual sequences in these clusters based on pairwise alignments to a reference database^48^ (**Fig. 1a and Supplementary Fig. 1a**). This reveals four broad clusters corresponding to the four main eukaryotic histones (H2A, H2B, H3, and H4) and their variants (H2A.Z, macroH2A, and cenH3), as well as a fifth cluster composed of archaeal HMfB homologs. Finally, this classification also uncovers three large connected components composed of transcription factors with histone-like DNA binding domains, which are widely distributed in eukaryotes (POLE3, POLE4, DR1) and/or archaea (NFYB). Further analysis of the genomic distribution of these histone genes shows a frequent occurrence of H3-H4 and H2A-H2B pairs in head-to-head orientation (5’ to 5’), strongly indicating co-regulation across eukaryotes (**Supplementary Fig. 1b,c and Supplementary Table 2**).

Next, we investigated the distribution and conservation of hPTMs across major eukaryotic groups and Archaea, including methylations, acetylations, crotonylations, phosphorylations, and ubiquitylations. To this end, histones from 19 different eukaryotic species were extracted, chemically derivatized^49^ and analyzed by mass-spectrometry (**Fig. 1c and Supplementary Table 3**), adding to previously available hPTM proteomics data for additional seven species. Our extensive taxon sampling covers all major eukaryotic groups, as well as hitherto unsampled early-diverging eukaryotic lineages—such as the malawimonad *Gefionella okellyi*, the discoban *Naegleria gruberi*, or the ancyromonad *Fabomonas tropica*—, thus providing a comprehensive comparative framework for evolutionary inference.

We focused first on hPTMs present in canonical histones, as defined by their highly conserved *N*-terminal regions, phylogenetic analyses, and sequence similarity to curated reference canonical histones (**Fig. 1d**; see Methods). hPTMs are detected in all canonical histones from all species. After correcting by sequence coverage, we observe that hPTMs are particularly abundant in H3 canonical histones (median = 23.5 hPTMs per species, mean = 24.3), compared with H2A, H2B and H4 (medians between 6.5 and 9, means between 9.5 and 13.4; **Supplementary Fig. 2a**). Holozoan canonical H2As (*Homo sapiens, Sycon ciliatum* and *Capsaspora owczarzaki*) represent an exception to this trend and contain similar number of modifications to H3s in these species. We also examined the reproducibility of hPTM detection across replicate samples, showing that the majority of hPTMs (87.5%) can be found in more than one sample (**Supplementary Fig. 2b,c**). Despite this, it is worth emphasizing that our data may contain false negatives, beyond the lack of coverage for particular residues that we systematically report. For example, some marks might be globally too scarce in the nucleosomes of a particular species, while other modifications like phosphorylations and ubiquitination are difficult to detect by mass-spectrometry without dedicated peptide-enrichment protocols.

Canonical H3 and H4 *N*-terminal tails contain the majority of phylogenetically-conserved hPTMs, in stark contrast with the relative paucity of conserved hPTMs in canonical H2A and H2B. A striking example of paneukaryotic conservation comes from the acetylation of the H4 K5, K8, K12 and K16 residues (**Fig. 1d**, second panel), all of which mark gene expression-permissive chromatin environments in multiple eukaryotic species^22^. A similar conservation pattern is observed in the acetylation of a group of *N*-terminal H3 lysines (K9, K14, K18, K23, K27) associated with similar functions, while other H3 acetylations are only found in a few species (e.g., residues K4, K56 and K79). While acetylations are highly conserved, only seven histone H3/H4 methylations are broadly conserved across eukaryotic lineages: H3K4me1/2/3, H3K9me1/2/3, H3K27me1/2/3, H3K36me1/2/3, H3K37me1/2/3 and, more sparsely, H3K79me1/2 and H4K20me1. Many of these broadly conserved marks have conserved roles in demarcating active (e.g., H3K4me) and repressive chromatin states (e.g., H3K9me and H3K27me)^22,42,50^. The scarcity of conserved hPTMs in H2A and H2B canonical histones can partially explained by their higher degree of sequence divergence (**Fig. 1e**), which is reflected in many non-homologous lysine residues (**Fig. 1d**). But even among homologous positions, we found little evidence of conservation, with the exception of H2A K5ac (associated to active promoters^51^) and, in fewer species, methylation of H2A K5 and H2B K5. Finally, we were also able to identify phosphorylations in serine and threonine residues and a few instances of ubiquitylation. In general, these marks show more restricted phylogenetic distributions than lysine acetylation or methylation, even in the tightly conserved H3 and H4 histones. We can identify conserved phosphorylations in H2A T120 and S122, which are shared by most opisthokonts, and the ubiquitylation of H2A K119 only in some holozoan species.

Mass-spectrometry analysis detected histone variants in all species included in our study, suggesting that they are relatively abundant in the chromatin of these eukaryotes (**Fig. 1e**). Most of these variants are lineage-specific, with the exception of the paneukaryotic variants H2A.Z, H3/cenH3 and H3.3; and the macroH2A variant found in holozoans and *Meteora* sp. (belonging to an orphan eukaryotic lineage). Interestingly, we find hPTMs in the vast majority of detected variants, both conserved and lineage-specific, particularly acetylations and methylations (**Fig. 1e and Supplementary Fig. 2d**). Overall, our comparative proteomic analysis suggests the existence of a highly conserved set of canonical hPTMs of ancestral eukaryotic origin in H3 and H4, which co-exists with less conserved hPTMs in H2A, H2B, and lineage-specific modifications in variant histones.

### Archaeal histones and histone post-translational modifications

In contrast with the paneukaryotic distribution of histones, sequence searches show that only a fraction of archaeal genomes encode for histones (28.1% of the taxa here examined; **Fig. 2a**). Archaeal histones exhibit a patchy phylogenetic distribution, similar to other gene families shared with eukaryotes^52^. Among others, histones are present in Euryarchaeota, the TACK superphylum and Asgard archaea^12,53–56^. Asgard are generally are considered to be the closest known archaeal relatives of eukaryotes^57,58^, although this sister-group relationship has been challenged by some studies^59^. Our extended sampling revealed that Asgard archaea histones, particularly in the Lokiarchaeota and Heimdallarchaeota clades^55^, often have lysine-rich *N*-terminal tails in the manner of eukaryotic histones (**Fig. 2a-c**). These Asgard histones appear to be conserved across multiple taxa, albeit without direct sequence similarity compared to canonical eukaryotic histones (**Supplementary Fig. 1d**). When compared against eukaryotic sequences classified in HistoneDB^48^, these archaeal histones clearly cluster in a separate group and are most similar to either eukaryotic H4 or, to a lesser degree, H3 canonical histones, in line with previous findings^12,55,60^.

**Figure 2.**
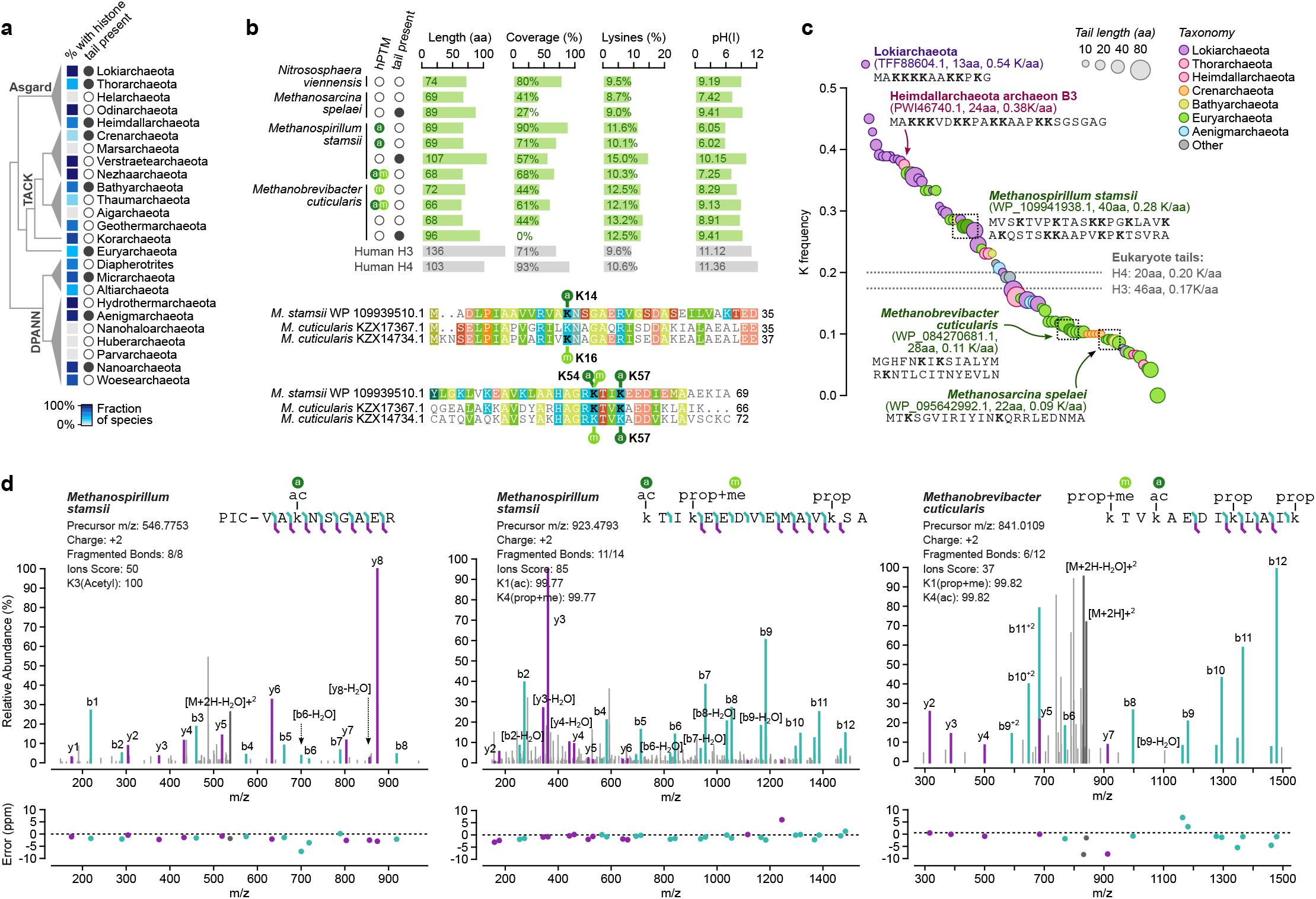
Archaeal histone diversity and post-translational modifications. **a,** Distribution of histones (fraction of taxa in each lineage) and histone tails (presence/absence) across Archaea phyla. **b,** Summary of proteomics evidence of archaeal histones, including the presence of modifications, tails, coverage, fraction of lysines identified, and isoelectric points. Human Histone H3 and H4 are included for reference. The alignments at the bottom depict the position of lysine modifications in the globular part of *Methanospirillum stamsii* and *Methanobrevibacter cuticularis* HMfB histones (modified residues in bold). **c,** Archaeal HMfB histones with *N*-terminal tails (at least 10 aa before a complete globular domain), sorted by frequency of lysine residues in the tail and color-coded according to taxonomy (same as panel A). Amino-acid sequences shown for selected examples. The dotted line indicates the median frequency of lysines in canonical eukaryotic H3 and H4 histone tails. Source data available in Supplementary Table 2. **d,** Mass spectra of three modified archaeal peptides, representing the relative abundance of fragments at various mass-to-charge ratios (m/z). Spectra were annotated using IPSA. b and y ions and their losses of H_2_O are marked in green and purple, respectively; precursor ions are marked in dark grey. Unassigned peaks are marked in light grey. Some labels have been omitted to facilitate readability.

To identify potential archaeal hPTMs, we performed proteomics analysis of histones in three Euryarchaeota (the Methanobacteriota *Methanobrevibacter cuticularis* and the Halobacteriota *Methanospirillum stamsii* and *Methanosarcina spelaei*) and one Thaumarchaeota species (*Nitrososphaera viennensis*; **Fig. 2b**). Mass-spectrometry detects histone proteins in all of them: 2-4 in the euryarchaeotes (with 27-90% protein coverage) and one in the thaumarchaeote (80% protein coverage), including homologs with *N*-terminal tails encoded by each of the three euryarchaeotes in our survey (22-40 aa, 0.09-28 lysines per residue; **Fig. 2c**). Moreover, this proteomics analysis finds evidence of hPTMs in archaeal histones. However, in comparison with eukaryotic histones, hPTMs are extremely scarce in archaeal histones. Specifically, we identify no hPTMs in *N. viennensis* and *M. spelaei* (one and two histones detected, respectively), three acetylations and one methylation in *M. stamsii* (in three out of four histones detected), and one acetylation and two methylations in *M. cuticularis* (in two out of four histones; **Fig. 2b,** top). Interestingly, we find conserved lysine residues with shared modifications in *M. stamsii* and *M. cuticularis* (methylation in K54 and acetylation in K57; **Fig. 2b,** bottom). This result indicates that highly-abundant hPTMs represent a eukaryotic innovation, likely linked to dynamic nucleosomal regulation in eukaryotes but not in Archaea.

### Taxonomic distribution of chromatin-associated proteins

hPTMs are deposited and removed by specific modifying enzymes (‘writers’ and ‘erasers’), while ‘reader’ protein domains found in diverse proteins bind and recognize specific hPTMs. For example, Bromo and Chromo domains bind acetylated and methylated lysine residues, respectively. In addition, the control of histone loading/eviction from specific genomic *loci* is mediated by chromatin remodellers, like SNF2 proteins^27^, and histone chaperones^26^. To date, the classification and evolutionary analysis of this chromatin machinery has been based on biased, partial taxonomic samplings and has not employed phylogenetic methods^61^ (with rare exceptions^12,27^), often resulting in inaccurate orthologous relationships and confounded classification and naming schemes.

We sought to obtain a systematic, phylogenetics-based classification of histone remodellers, chaperones, readers, and modifiers in order to understand the evolutionary history of eukaryotic chromatin (**Fig. 3a**). To this end, we (*i*) compiled a taxa-rich dataset of 172 eukaryotic genomes and transcriptomes, covering all major eukaryotic supergroups and devoting particular attention to early-branching, non-parasitic lineages (**Supplementary Table 1**), as well as genomic data from 4,226 Archaea, 24,886 Bacteria and 185,579 viral taxa; (*ii*) defined a protein structural domain as a proxy for each gene family (**Supplementary Table 4**) and retrieved all genes in these genomes that contained these domains; and (*iii*) inferred accurate orthology groups from phylogenetic analyses of each gene class (next section).

**Figure 3.**
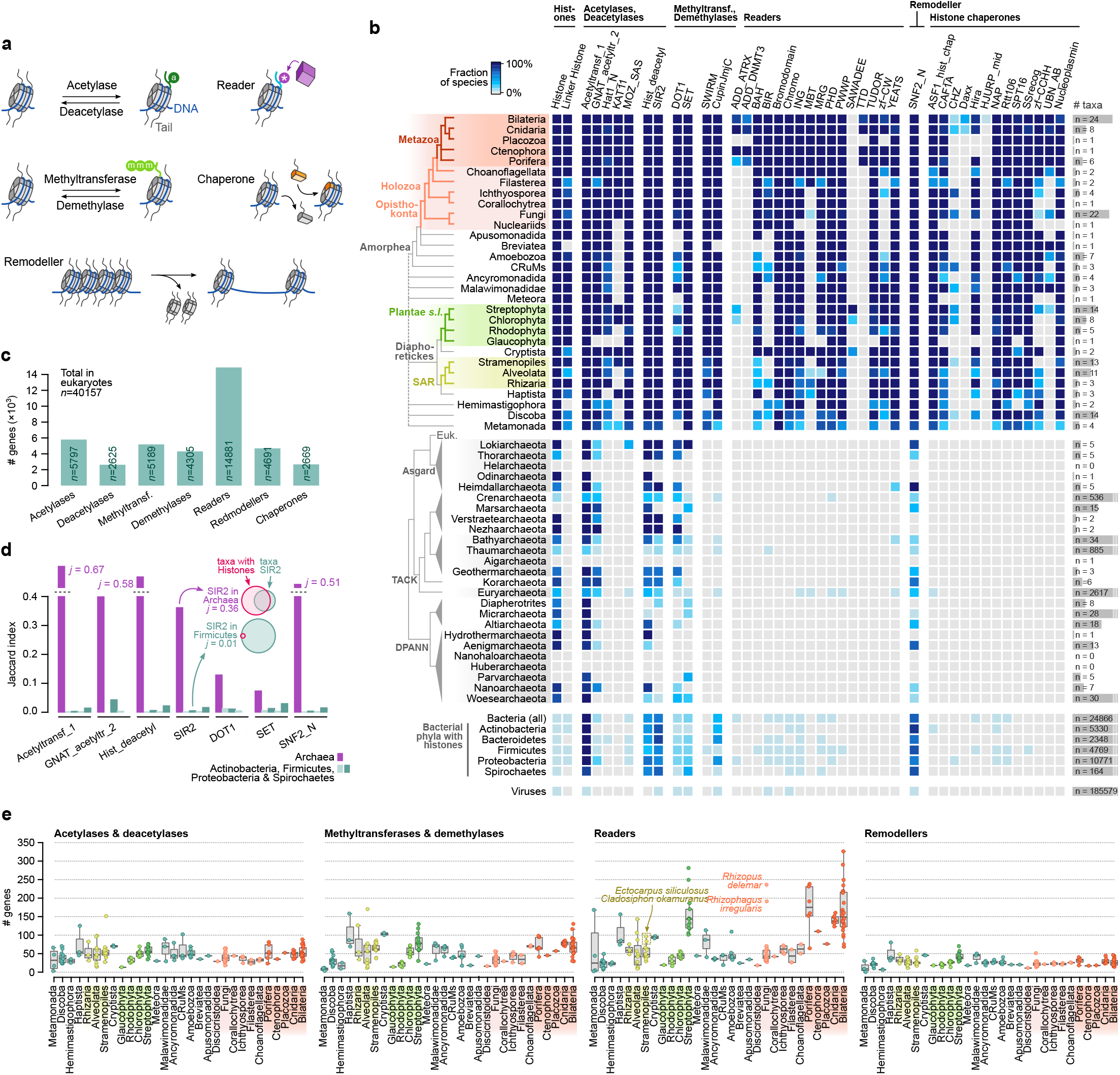
Taxonomic distribution of chromatin-associated gene classes. **a,** Summary of the seven classes of genes with chromatin-related activity covered in our survey: histone-specific hPTM writers (acetylases and methyltransferases), erasers (deacetylases and demethylases), readers, remodellers, and chaperones. **b,** Percentage of surveyed taxa containing homologs from each chromatin-associated gene class, for eukaryotes (top), archaea, bacteria, and viruses (bottom). Species-level tables are available in Supplementary Fig. 3. **c,** Number of eukaryotic genes classified in each of the chromatin-associated modification enzymes, readers, remodellers, and chaperones**. d,** Overlap between the taxon-level phylogenetic distribution of histones and chromatin-associated domains in archaea and four bacterial phyla, measured using the Jaccard index. **e,** Number of genes encoding writer, eraser, reader and remodeller domains, per species.

We examined the taxonomic distribution and abundance of the major gene classes (**Fig. 3b,c**). Many domains with chromatin-associated functions in eukaryotes are also present in Archaea and Bacteria, albeit with scattered phylogenetic distributions (**Fig. 3b and Supplementary Fig. 3a,b**). Families with prokaryotic homologs include mostly catalytic gene classes (writer, eraser and remodeller enzymes), whereas readers and histone chaperones are virtually absent from prokaryotes (**Fig. 3b**). Histone fold-encoding genes constitute a case in point for this patchy distribution of chromatin proteins in prokaryotes: they are present in most archaeal phyla, but are absent in about half of the sampled genomes within each (**Fig. 3b**). Yet, there is a qualitative difference between the phylogenetic distribution of archaeal and bacterial chromatin-associated gene classes: whereas archaeal histones tend to co-occur with chromatin-associated gene classes, the bacterial complement of writers and erasers is much less conserved and is uncorrelated with the extremely rare presence of histone-like genes (**Fig. 3d**).

Within eukaryotes, most gene structural classes associated with chromatin functions are ubiquitously distributed across all lineages here surveyed, supporting an early eukaryotic origin for the core chromatin machinery (**Fig. 3b and Supplementary Fig. 3d**). In fact, the total number of chromatin writer, eraser and remodeller enzymes remains remarkably stable across eukaryotes (**Fig. 3e**). The only exception is the marked increase in genes encoding reader domains observed in lineages exhibiting complex multicellularity: animals, streptophyte plants, and, to a lesser degree, phaeophyte brown algae (Stramenopila). This occurs partially due to the addition of new gene classes (e.g., SAWADEE in the Plantae *s.l*. + Cryptista lineage, or ADD_DNMT3 in bilaterians and cnidarians), but also via the expansion of ancient, widely-distributed reader gene classes (e.g., Tudor, PHD, Chromo or Bromo domains). These taxonomic patterns indicate that chromatin modifying and remodelling catalytic activities originated in prokaryotes, while reader and chaperone structural domains are eukaryotic innovations.

### Phylogenetics of chromatin modifiers and remodellers

To gain detailed insights into the origin and evolution of chromatin gene families, we used phylogenetic analysis to define orthology groups from paneukaryotic gene trees. We surveyed 172 eukaryotic species and defined a total of 1,713 gene families (orthogroups) encompassing 51,426 genes, 95% of which were conserved in two or more high-ranking taxonomic groups (as listed in **Fig. 1a**), and which included 51,426 genes in total (**Supplementary Table 5**). We annotated each gene family according to known members from eukaryotic model species. For simplicity, we use a human-based naming scheme throughout the present manuscript (unless otherwise stated), but we also provide a dictionary of orthologs in three additional model species (*Arabidopsis thaliana, Saccharomyces cerevisiae* and *Drosophila melanogaster*; see **Supplementary Table 5**). This phylogenetic classification scheme of eukaryotic chromatin gene families, as well as the sequences and associated phylogenetic trees, can be explored and retrieved in an interactive database: https://sebe-lab.shinyapps.io/chromatin_evolution

We first investigated the potential pre-eukaryotic origins of these gene families/orthogroups by comparing their phylogenetic distance to prokaryotic sequences and to other eukaryotic orthogroups (**Fig. 4a**). Most eukaryotic gene families are more closely related to other eukaryotes than to prokaryotic sequences, supporting the idea that writers, erasers, remodellers and readers diversified within the eukaryotic lineage, as previously noted for histones^12^. This analysis also reveals a substantial fraction of eukaryotic gene families with close orthogroups in Archaea and Bacteria, which pinpoints components that were (*i*) inherited from a prokaryotic ancestor during eukaryogenesis; (*ii*) laterally transferred between eukaryotes and prokaryotes at later stages; or (*iii*) a combination of both phenomena. For example, we identified a well-supported sister-group relationship between the eukaryotic SIRT7 deacetylase and a clade of Asgard archaea Sirtuin enzymes (Heimdallarchaeota and Lokiarchaeota), a topology compatible with an archaeal origin or ancient transfers to/from Asgard and eukaryotes^62^; whereas SIRT6 appears nested within other eukaryotic sequences (**Fig. 4b**, left). Likewise, the KAT14 acetylase is more closely related to bacterial enzymes than to other eukaryotic acetylases (**Fig. 4b**, right).

**Figure 4.**
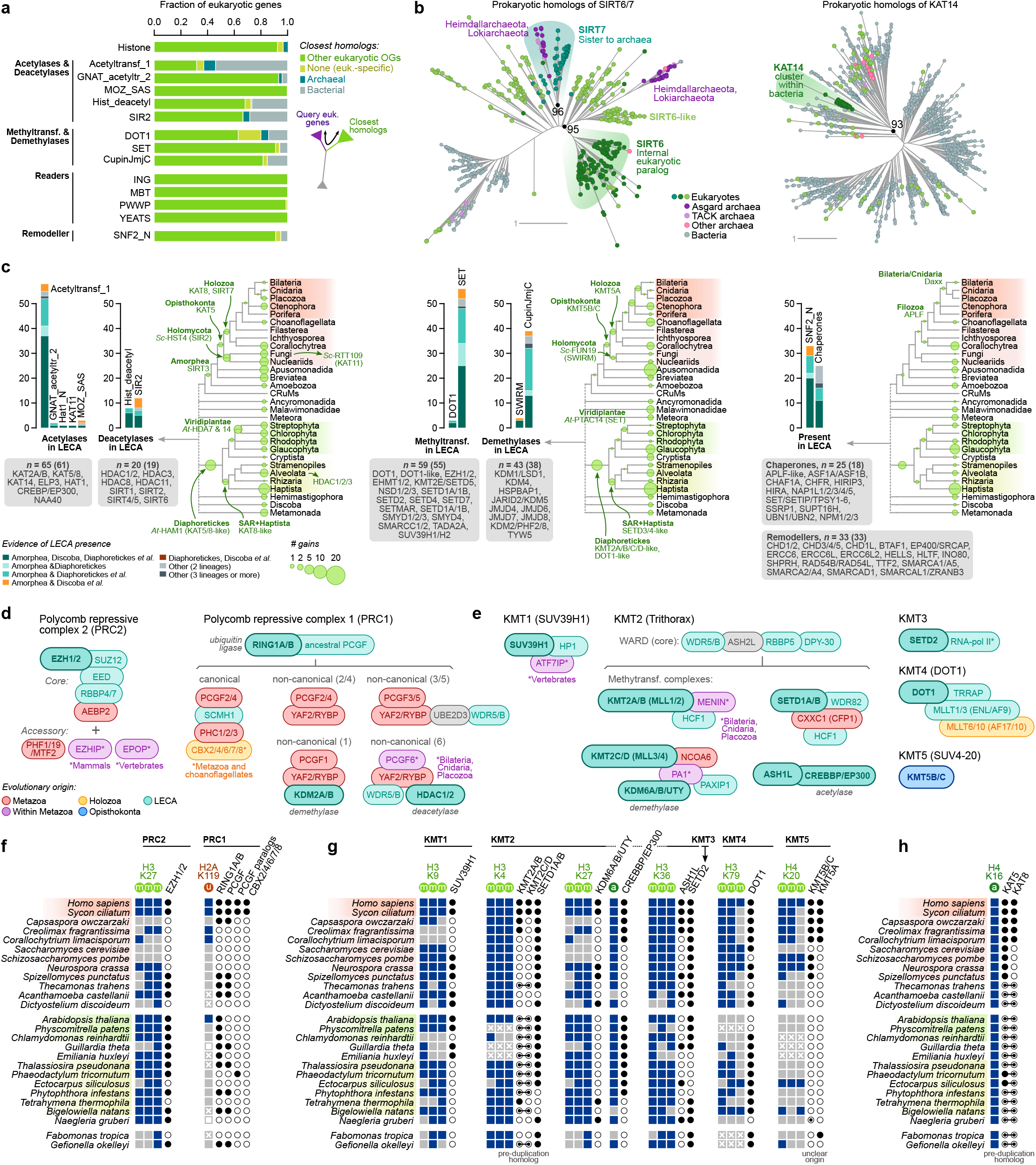
Origin and evolution of chromatin-associated gene families. **a,** Summary of phylogenetic affinities of the eukaryotic homologs of gene classes that are also present in prokaryotes. For each gene family, we evaluate whether it is phylogenetically closer to a majority (≥50%) of eukaryotic sequences from a different orthogroup (indicating intra-eukaryotic diversification), or to sequences from Bacteria or Archaea. **b,** Left, gene tree of eukaryotic and prokaryotic Sirtuin deacetylases, showcasing an example of a eukaryotic family that diversified within eukaryotes (SIRT6) and another one with close relatives in Asgard archaea (SIRT7). Right, gene tree of KAT14 acetylase, a eukaryotic orthogroup with bacterial origins. Statistical supports (UF bootstrap) are shown at selected internal nodes of the highlighted clades. **c,** Evolutionary reconstruction of hPTM writer and eraser gene families, remodellers, and histone chaperones along the eukaryotic phylogeny, including the number of genes present in the last eukaryotic common ancestor (LECA). Barplots indicate the number of orthologs of each gene family present at the LECA (at 90% posterior probability; see Methods) and whether the presence of a given orthogroup at LECA is supported by its conservation in various early-branching eukaryotic lineages (Amorphea, Discoba, Diaphoretickes and others). The list of ancestral gene families below each plot is non-exhaustive. Two ancestral gene counts are provided: all families at presence probability above 90%, and, in brackets, the subset of these that is present in at least two of the main eukaryotic early-branching lineages (Amorphea, Diaphoretickes, and Discoba). Source data in Supplementary Table 5. **d-e,** Reconstructed evolutionary origins of the different subunits of the Polycomb repressive complexes (PRC2 and PRC1) and Trithorax-group complexes (KMT1 to 5). **f-h,** Side-by-side comparison of the presence of individual hPTM marks and various subunits of the Polycomb and Trithorax complexes, as well as other hPTM writers, responsible for their deposition.

Next, we mapped the phylogenetic distribution of orthogroups in order to infer the origin and diversification of individual chromatin gene families (**Fig. 4c and Supplementary Fig. 4a**). Using probabilistic inference of ancestral gene content, we reconstruct a rich Last Eukaryotic Common Ancestor (LECA) complement of chromatin-associated gene families: 65 acetylases (amongst which 61 were conserved in at least two of the most deeply sampled eukaryotic early-branching lineages, namely Amorphea, Diaphoretickes, and Discoba); 20 deacetylases (19 in these early-branching eukaryotic lineages); 59 methyltransferases (55); 42 demethylases (38); 33 remodellers (33); and 25 chaperones (18) (**Fig. 4c and Supplementary Table 5**). The subsequent evolution of these families is characterized by relative stasis, with few new orthologous families emerging in later-branching eukaryotic lineages. Notable exceptions include the origin of KAT5 deacetylases and KMT5B/C SET methyltransferases in Opisthokonta; KAT8 and SIRT7 in Holozoa; and Viridiplantae-specific deacetylases (homologs of *A. thaliana* HDA7 and HDA14 deacetylases) and SETs (*A. thaliana* PTAC14); among others.

In spite of their broad distributions across eukaryotes, many chromatin modifier families exhibit variation in their protein domain architectures, likely conferring them functional properties such as distinct binding preferences (**Supplementary Fig. 4b**). For example, most CREBBP/EP300 acetylases consist of a catalytic HAT_KAT11 domain and two TAZ and ZZ zinc finger domains, but different lineages have acquired different reader domains: an acetylation-reading Bromo domain in holozoans and stramenopiles, PHD in plants and some stramenopiles, and no known reader domains in other lineages (e.g., in the fungal orthologs of the *S. cerevisiae* protein RTT109). A similar pattern is apparent in SET methyltransferase families sharing a core catalytic domain (SET) harboring variable DNA- and chromatin-interacting domains – animal SETDB1/2 homologs have MBD domains that bind CpG methylated DNA, while plants have SAD_SAR domains with the same function; and holozoan ASH1L homologs encode Bromo and BAH readers, whereas phaeophytes encode PHD domains (**Supplementary Fig. 4b**). Other architectures, however, are much more conserved, as exemplified by the presence of Tudor-knot and MYST zinc finger domains in most KAT5 deacetylases; or the ubiquitous co-occurrence of Helicase-C and SNF2_N domains in most remodellers (**Supplementary Fig. 4b**).

Specific examples of evolutionarily conserved chromatin gene families include the catalytic core and the subunits of well-studied chromatin complexes^63^ like PRC1 (RING1/AB, PCGF), PRC2 (EZH1/2, SUZ12, EED, RBBP4/7) and Trithorax/MLL (MLL1/2/3/4, WRD5, ASH2L, RBBP5, DPY-30; **Fig. 4d,e**). However, when we compared the distribution of these complexes with the hPTMs they are related to, we found a generally poor co-occurrence (**Fig. 4f-h**). For example, organisms like *Dictyostelium discoideum* and *Creolimax fragrantissima* lack EZH1/2 orthologs, but we detected H3K27me3 in these species; while *Thecamonas trahens* and *Naegleria gruberi* lack Dot1 orthologs but have H3K79me marks. A poor correlation is also observed between the occurrence of H3K9me and that of SUV39H1 orthologs. An exception to this pattern is the ubiquitous distribution of H4K16ac and the acetylase family KAT5/8^64^ (**Fig. 4h**). These patterns suggest that the specificity between hPTMs and their writers might not be completely conserved across eukaryotes, with distinct members of the same gene classes (e.g., methyltransferases) performing similar roles. In this context, reading domains present in writing/erasing enzymes (directly in the same protein or as part of multi-protein complexes) are likely to play a major role in the re-purposing of chromatin catalytic activities.

### Evolutionary expansion of chromatin readers

Multiple protein structural domains have been involved in the recognition of hPTMs, such as Bromo and PHD domains binding to acetylated lysines or Chromo, MBT and Tudor domains binding to methylated lysines^23,24^. These are generally small domains and can be found both as stand-alone proteins as well as in combination with other domains, often catalytic activities such as hPTM writers, erasers and remodellers. Thus, they are central in the establishment of functional connections between chromatin states. To understand the contribution of these reading domains to the evolutionary diversification of chromatin networks, we studied in detail the phylogeny and protein architecture of reader domains across eukaryotes.

We quantified the co-occurrence frequency of reader and catalytic domains, finding (*i*) that most reader domains are present in genes without writer, eraser or remodeller domains (87%, **Fig. 5a**); and (*ii*) that most cases of reader-catalytic co-occurrence involve PHD, Chromo and Bromo domains (**Supplementary Fig. 5a**). For example, the conserved architecture of the paneukaryotic CHD3/4/5 re-modellers includes Chromo readers in most species and PHD domains specifically in animals and plants (**Supplementary Fig. 4b**). Likewise, PHD domains are often present in the KMT2A/B and KMT2C/D SET methyltransfrase; and the ASH1L family has recruited Bromo and BAH domains in holozoans, and PHD in multicellular stramenopiles (**Supplementary Fig. 4b**). In spite of these redundancies, reader families typically have independent evolutionary histories, as illustrated by the fact that most reader domain-containing genes encode only one such domain (92%, **Supplementary Fig. 5b**).

**Figure 5.**
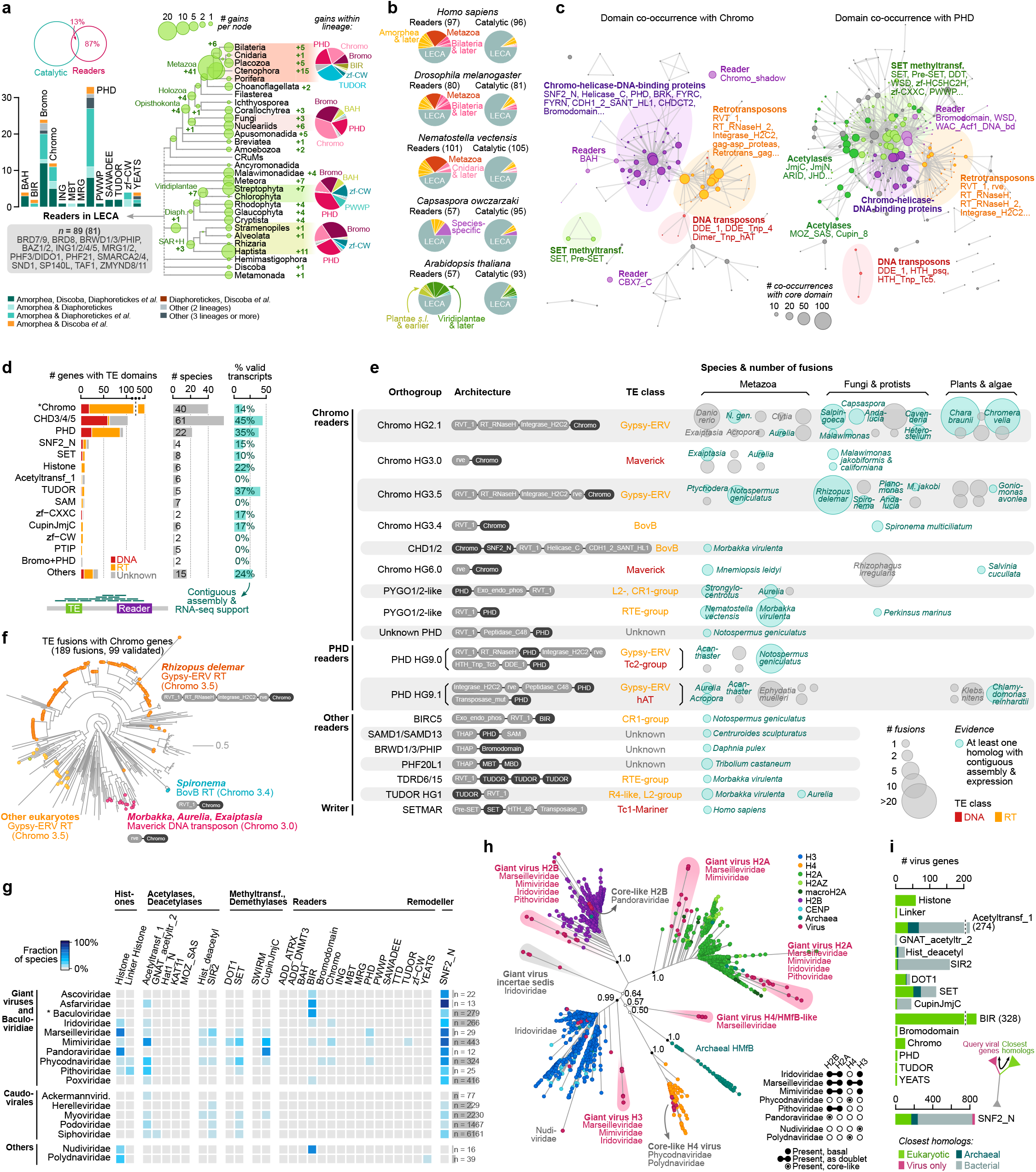
Evolution of chromatin readers and capture of chromatin proteins by transposable elements and viruses. **a,** Evolutionary reconstruction of reader gene families along the eukaryotic phylogeny, highlighting the number of gains along the eukaryotic phylogeny (at 90% posterior probability). The Euler diagram at the top shows the overlap between presence of chromatin-associated catalytic domains and readers. The barplot at the left indicates the number of orthologs of each gene family present at the LECA and whether their presence is supported by its conservation in various early-branching eukaryotic lineages (Amorphea, Discoba, Diaphoretickes, and others). Pie plots at the right summarize the number of orthogroups from each gene family gained within selected lineages: Metazoa, Holomycota, Viridiplantae and SAR+Haptophyta. **b,** Number of reader or catalytic orthogroups gained at each node in the species tree, for selected species. Source data in Supplementary Table 5. **c,** Networks of protein domain co-occurrence for Chromo and PHD readers. Each node represents a protein domain that co-occurs with Chromo or PHD domains, and node size denotes the number of co-occurrences with either Chromo or PHD. Edges represent co-occurrences between domains. Groups of frequently co-occurring protein domains have been manually annotated and color-coded, which has revealed sub-sets of retrotransposon and DNA transposon-associated domains. **d,** Number of chromatin-related eukaryotic genes fused with transposons grouped by gene family (left), including the fraction that are classified as valid gene models based on expression and assembly data (centre); and the number of species where each type of fusion is found (right). The number of fusion events are colored according to their similarity with known DNA transposons (red) or retrotransposons (orange) from the Dfam database (see Methods). (*) The ‘Chromo’ category excludes genes containing other chromatin-associated protein domains such as SNF2_N (listed separately as ‘Chromo+SNF2_N’, which includes remodellers with the domain of unknown function DUF1087, which is also common in DNA transposons). **e,** Selected examples of transposon fusion domains classified by orthogroup, including their archetypical protein domain architecture, homology to transposon class, their phylogenetic distribution, and number of fusion genes. Only orthogroups with at least one valid gene model are listed. Source data available in Supplementary Table 6. **f,** Example tree of Chromo readers, highlighting genes with fused TE-associated domains and their consensus domain architectures. **g,** Fraction of viral genomes containing homologs from each chromatin gene family, for nucleocytoplasmic giant DNA virus families (top) and other taxa containing histone domains (Nudiviridae, Polydnaviridae; bottom). **h,** Phylogenetic analysis of histone domains, with a focus on viral homologs. Statistical supports (approximate Bayes posterior probabilities) are shown for the deepest node of each canonical eukaryotic or archaeal histone clade. The inset table summarizes the presence of doublet histone genes per linage. **i,** Number of viral homologs in each chromatin-associated gene family, classified according to their closest cellular homologs (eukaryotes, bacteria or archaea) in phylogenetic analyses (see Methods). Source data available in Supplementary Table 6.

We next performed phylogenetic analyses of individual reader domains and reconstructed the gains and losses of these reader gene families/orthogroups (**Fig. 5a**). Compared to the relative stasis of catalytic enzyme families, this reader-centric analysis revealed a strikingly different evolutionary pattern of lineage-specific bursts of innovation, particularly amongst PHD, Chromo and Bromo genes, as well as Tudor in animals (**Fig. 5a and Supplementary Fig. 5c**). PHD, Chromo and Bromo families also appeared as the most abundant in the reconstructed LECA reader domain repertoire, which amounted to 89 gene families (**Fig. 5a**, left). The distribution of gene family ages in extant species also corroborates that more readers have emerged at evolutionarily more recent nodes of the tree of life than catalytic gene families (**Fig. 5b**).

### Co-option of the chromatin machinery by transposable elements

Further examination of the domain co-occurrence networks of readers revealed that Chromo and PHD domains are often present together with protein domains found in transposable elements (TEs; **Fig. 5c and Supplementary Table 6**), including retrotransposons (e.g., retrotranscriptases and integrases; orange modules in **Fig. 5c**) and DNA transposons (e.g., DNA binding domains and transposases; red modules). It is known that some TEs show insertion-preferences associated to specific chromatin states^65^, often mediated by direct chromatin tethering mechanisms^66^. For example, the Chromo domain of the MAGGY gypsy retrotransposon of the fungus *Magnaporthe grisea* targets H3K9me regions^67^. Reciprocally, some protein domains of TE origin, often DNA-binding domains, have been co-opted into chromatin and transcriptional regulators^68^. Thus, we decided to explore in detail the occurrence of chromatin-associated domain (readers, but also catalytic domains) linked to TEs in the 172 eukaryotic genomes in our dataset (**Fig. 5d**). Moreover, we used available RNA-seq datasets in many of these species to validate some of these TE fusions (**Fig. 5d-e**). A fully validated fusion gene would (*i*) come from a non-discontinuous gene model in the original assembly, and (*ii*) have evidence of expression, with reads mapping along the entire region between the TE-associated domain and the chromatin-associated domain (**Supplementary Fig. S6**).

We identified 823 predicted gene models containing both chromatin- and TE-associated domains (**Fig. 5d**). Whilst these TE fusions were not exclusive of reader domains, most such fusions involved PHD and Chromo-encoding genes; followed by SNF2_N remodellers, SET methyltransferases, and others. An homology search against a database of eukaryotic TEs revealed that most of these candidate TE fusions could be aligned to known retrotransposons or DNA transposons. For example, by way of validation, our analysis identifies the SETMAR human gene, a previously-described fusion between a SET methyltranferase and a Mariner-class DNA transposon^69^. Overall, 31% of the candidate fusion genes were supported by valid gene models according to our stringent criteria (**Fig. 5d**). Interestingly, we find very few cases of hypothetical fusions between TEs and Bromo domains, which recognize K acetylations and are otherwise highly abundant across eukaryotes, and none of them is validated by RNA-seq data. This could be explained by the detrimental effect of targeting TE insertions to sites of active chromatin demarcated by histone acetylations, such as promoter and enhancer elements.

Some of these validated fusions have a broad phylogenetic distribution (**Fig. 5e**), such as a Gypsy-ERV retrotransposon with a *C*-terminal Chromo domain (Unk. Chromo 2.1 in **Fig. 5e**) that is widely distributed in animals and various microbial eukaryotes, and contains dozens of paralogs in vertebrate *Danio rerio* or the charophyte *Chara braunii*, many of which are expressed. Another widespread Gypsy-ERV retrotransposon with a Chromo domain is present in multiple expressed and highly similar copies in the fungus *Rhizopus delemar* (**Fig. 5f,e**), suggesting a successful colonization of this genome by this TE. By contrast, other TE fusions are taxonomically restricted to one or few related species, such as the fusion of hAT activator DNA transposons with Chromo CBX and CDY readers in the sponge *Ephydatia muelleri*; or multiple instances of fusions with Chromo and PHD readers in cnidarians. A common fusion in cnidarians involves different retrotransposon classes with PHD domains orthologous to the PYGO1/2 protein (**Fig. 5e**), which is known to recognize specifically H3K4me^70^. Globally, this analysis reveals that recruitment of chromatin reading and even modifying domains by TE has occurred in many eukaryotic species, in a way that might facilitate the evasion from suppressing mechanisms in the host genomes as suggested by the expansion of Chromo-fused TEs in the genomes of *Chara braunii* (Viridiplantae), *Chromera velia* (Alveolata) and *Rhizopus delemar* (Fungi).

### Chromatin components in viral genomes

In addition to TEs, chromatin is also involved in the suppression of another type of genomic parasites: viruses. Some chromatin-related genes, including histones, have been found in viral genomes, especially among the nucleocytoplasmic large DNA viruses – also known as giant viruses. Eukaryotic core histones have been even hypothesized to have evolved from giant virus homologs, after the discovery that certain Marseilleviridae genomes encoded deeply-diverging orthologs of the four canonical histones^71^. These viral histones have been recently shown to form nucleosome-like particles that package viral DNA^72,73^.

We analyzed the distribution and abundance of chromatin-related protein domains among viruses, including data from 1,816 giant virus genomes. Based on structural domain searches, we identified 2,163 viral chromatin-related proteins (**Fig. 5g and Supplementary Table 6**). The majority of these proteins are encoded by giant viruses (55%), followed by Caudovirales (37%). Among these two groups, only giant virus genomes encode histones – specifically, the Iridoviridae, Marseilleviridae, Mimiviridae, Pithoviridae, and Phycodnaviridae families. Concordantly with previous studies^74^, we also identify remodellers in all giant virus families; as well as less abundant components of the chromatin writer/eraser/reader toolkit (**Fig. 5g**).

We then investigated the phylogenetic affinities of these viral chromatin proteins, starting with histones (**Fig. 5h**). Our analysis recovers the phylogenetic affinity of Marseilleviridae histones with specific eukaryotic histone families^71^, and makes this pattern extensive to Mimiviridae, Iridoviridae, and Pithoviridae giant viruses (**Fig. 5h**), with the caveat of the ambiguous clustering of the H4-like viral histones with either H4 eukaryotic or archaeal HMfB genes. In all these lineages, we identify genes encoding two histone-fold domains orthologous to H2B + H2A (inset table in **Fig. 5h**), whereas the H4 + H3 histone doublet genes appears to be exclusive to Marseilleviridae. By contrast, histone homologs in Phycodnaviridae, Pandoraviridae (also giant viruses), and Polydnaviridae (*incertae sedis*) are never found as either doublets or as early-branching homologs of eukaryotic histones, suggesting recent acquisition from eukaryotes.

Unlike histones, most of the viral chromatin-associated genes exhibited a mixture of prokaryotic and eukaryotic phylogenetic affinities and often lack affinity to any specific eukaryotic gene family (**Fig. 5i and Supplementary Fig. 7**). Viral readers, on the other hand, are often embedded within eukaryotic clades in gene trees and are similar to *bona fide* eukaryotic families, exhibiting topologies consistent with recent, secondary acquisitions. This is the case of BIRC2/3/XIAP readers widespread in the Baculoviridae, which encode BIR domains that are often hijacked from their hosts^75^. We also find a number of viral Chromo-encoding genes, which fall in two main taxonomic categories: (*i*) giant virus homologs of the eukaryotic CBX1/3/5 family (present in Mimiviridae, Iridoviridae and Phycodnaviridae); and (*ii*) homologs from various Adintoviridae, which are closely related to animal Chromo genes encoding *rve* integrase domains^76^ (**Fig. 5i**). Finally, we also identify a handful of eukaryotic-like viral genes with deep-branching positions relative to core eukaryotic gene families, as seen in histones (**Fig. 5h**). This includes Mimiviridae homologs of the eukaryotic methyltransferases SMYD1-5 and DOT1 (**Supplementary Fig. 7d,e**), as well as SNF remodeller families with homologs in distinct giant virus clades (HLTF/TTF2 in Phycodnaviridae, Mimiviridae and Iridoviridae). These results indicate that cases of horizontal transfer from eukaryotes to viruses are common in different chromatin-related gene families, including histones. Therefore, it is likely that basally-branching giant virus histones were similarly acquired from a stem eukaryotic lineage and this would explain the observed histone tree topology with extant eukaryotic species. In any case, most of the eukaryotic chromatin machinery appears to have cellular roots.

## Discussion

Our comparative proteogenomics study reconstructs in detail the origin and evolutionary diversification of eukaryotic chromatin components, from post-translational modifications to gene family domain architectures. We looked first at the pre-eukaryotic roots of chromatin. Multiple aspects of archaeal chromatin have been studied in recent years, including nucleosomal patterns^31^ and the structure of the archaeal nucleosome^30^. A recent taxonomic survey of archaeal nucleoid-associated proteins revealed multiple independent diversifications of DNA-wrapping proteins and a strong association between high levels of chromatinization and growth temperature, overall suggesting a structural, non-regulatory role for archaeal chromatin^77^. Our proteomics data support this notion by showing the scarcity of hPTMs in four species belonging to two different archaeal lineages (Euryarchaeota and Thaumarchaeota). An earlier proteomics study reported the complete absence of hPTMs in the euryarchaeote *Methanococcus jannaschii*^34^. Here we do identify a few instances of modified lysine residues in Euryarchaeota, which is in line with the recently reported acetylations in *Thermococcus gammatolerans* histones^78^. It remains to be seen if hPTMs are frequently present in Asgard and other unsampled archaeal linages, where other eukaryotic-like features have been found^57,79,80^. In fact, some of these Asgard, particularly Lokiarchaeota, encode for histones with long, K-rich *N*-terminal tails but that bear no similarity with eukaryotic histones and are, therefore, most probably the result of convergent evolution. Interestingly, Lokiarchaeota genomes also frequently encode histone modifiers such as SET methyltransferases and MOZ_SAS acetylases. However, overall our results suggest that extensive usage of hPTMs is an eukaryotic innovation (**Fig. 6a**). Similarly, while we find the majority of catalytic domains of hPTM writers, hPTM erasers and chromatin remodellers in Archaea and even Bacteria, these appear only scattered in a small fraction of the examined taxa. In contrast, hPTM reader domains and histone chaperones are eukaryotic innovations, further supporting the idea that the functional readout of hPTMs and the role for histone variants in defining chromatin states are both exclusive to eukaryotes (**Fig. 6a**).

**Figure 6.**
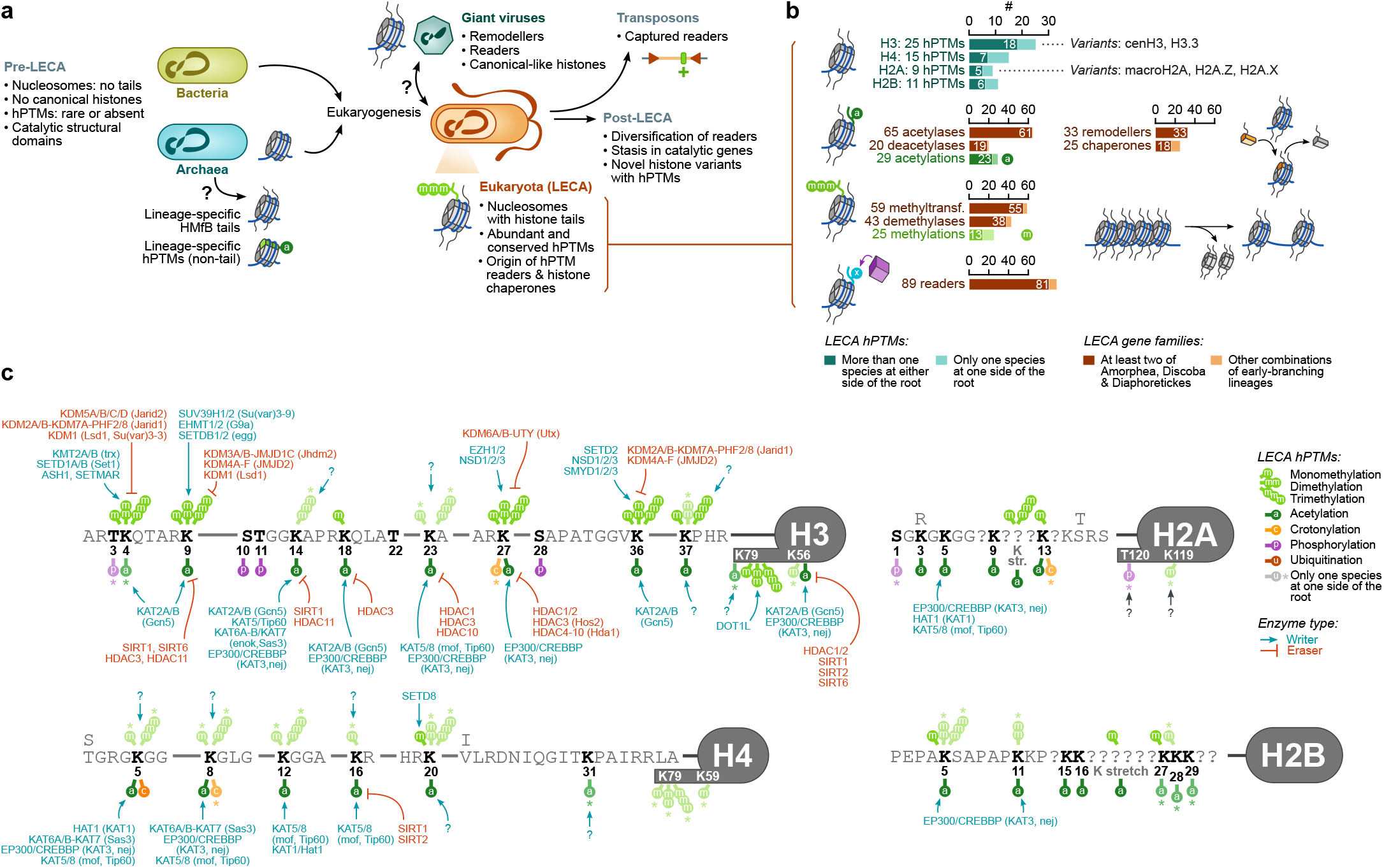
Chromatin evolution and eukaryogenesis. **a,** Summary of events in chromatin evolution prior to, during and after the origin of eukaryotes. **b,** Number of chromatin-related gene families and hPTM marks inferred to have been present at the LECA. Ancestral gene counts are indicated at >90% probability. For gene counts, numbers within bars indicate the subset of families present in at least two of the most deeply-sampled early-branching eukaryotic lineages (Amoropha, Diaphoretickes, and Discoba). For hPTMs, the ancestral counts have been inferred using Dollo parsimony assuming a Diaphoratickes – Amorphea split at the root of eukaryotes, and numbers within bars indicate the number of hPTMs whose ancestral presence is supported by more than one species at both sides of the root. **c,** hPTMs inferred to be present in the last eukaryotic common ancestor (LECA) based on Dollo parsimony. Only amino-acid positions conserved in all eukaryotes in our dataset are shown. Asterisks indicate modifications whose presence at the LECA is supported by just one species at either side of the root. The inferred LECA presence of known writing/erasing enzymes associated to these hPTM is indicated.

The origin of eukaryotes represents a major evolutionary transition in the history of life^81^. Thanks to sequencing and comparative analysis of archaeal and eukaryotic genomes, we also have a detailed reconstruction of the massive innovation in gene repertoires that occurred at the origin of eukaryotes. This gene innovation in the Last Eukaryotic Common Ancestor (LECA) includes cytoskeletal proteins and associated motors like myosins^82,83^ and kinesins^84^, vesicle trafficking apparatus^85^, splicing machinery^86^, ubiquitin signalling systems^87^ and a large repertoire of sequence-specific transcription factors^37^. Combining parsimony analysis and knowledge on gene function in extant lineages (mostly vertebrates, yeast and plants), our results allow us to reconstruct a complex LECA repertoire of hPTMs and associated writing, eraser and reader gene families (**Fig. 6b,c**). We infer 23 to 29 highly-conserved lysine acetylations in canonical histones (e.g., H3K9ac and H3K27ac) and a repertoire of 65 and 20 histone acetylase and deacetylase families, respectively. With the exception of H4K16ac^64^, most histone acetylations are thought to exert a generic, perhaps additive, effect on the opening of chromatin^22^. As such, acetylation marks like H3K27ac have been found to be enriched in promoters of active genes in diverse eukaryotes^42^. In contrast, histone methylations often have very specific readouts and they can be linked both to active and repressive chromatin states. We infer between 13 and 25 conserved methylated lysine residues in LECA histones, including marks typically associated to active promoters (H3K4me1/me2/me3), gene bodies (H3K36me3, H3K79me1/2, H4K20me1), and repressive chromatin states (H3K9me2/me3, H3K27me3, H4K20me3)^88,89^. Finally, we also infer the existence of five histone variants in the LECA (cenH3, H3.3, H2A.Z, macroH2A and H2A.X), as well 33 chromatin remodellers (e.g., EP400/SWR1 and INO80, involved in loading and removal of H2A.Z, respectively) and 25 histone chaperones (e.g., ASF1A/B and NPM1/2/3). This indicates that, in addition to an extensive repertoire of hPTMs, the regulation of nucleosomal histone composition was also an important feature in the LECA.

Chromatin evolution after the origin of eukaryotes is characterized by an expansion of lineage-specific histone variants harboring unique hPTMs and a net expansion in the number of reader gene families, as opposed to the relatively static catalytic gene families (writers, erasers and remodellers). This is particularly relevant as it suggests extensive remodelling of chromatin networks during eukaryote evolution, that is, changes in the coupling of particular hPTMs to specific functional chromatin states. An example of such changing state-definitions comes from looking at the hPTMs associated to TEs in different organisms: H3K9me3+H4K20me3 in animals, H3K27me3 in some plants^90^, H3K79me2+H4K20me3 in the brown multicellular algae *Ectocarpus siliculosus*^43^, and H3K9me3+H3K27me3 in the ciliate *Paramecium tetraurelia*^91^. In the context of the histone code hypothesis^3,20,92–94^, our findings indicate that, while there is an ancient core of conserved hPTMs across eukaryotes, evidence for a universal code/functional-readout is limited, with perhaps the exception of the highly conserved configuration of ancient hPTMs around active promoters across many eukaryotes^42^. Another interesting observation related to the evolution of chromatin networks is the capture of chromatin reader domains by TEs. We find evidence of this phenomenon in a number of species with a scattered phylogenetic distribution, suggesting that it is a recurrent process and that it often leads to the successful propagation of the TE in the host genome. We hypothesize that this process facilitates the targeting of TEs to specific chromatin states, as it has been described in the case of MBD DNA methylation readers captured by TEs^95,96^.

In the future, a broader phylogenetic understanding of the genome-wide distribution of hPTMs, as well as the direct interrogation of hPTM binders in different species^97–99^, will be crucial to further clarify questions such as the ancestral role of specific hPTM and the co-option of ancient hPTMs into novel functions.

## Acknowledgements

We want to thank Alex de Mendoza for critical input on the analysis of transposable element fusions. We also want to thank Josep Casacuberta for *Physcomitrella patens* samples, Harold J. G. Meijer for *Phytophthora infestans* samples, Maja Adamska for *Sycon ciliatum* samples, and Alistar Simpson for access to the *Gefionella okellyi* culture (made possible by his funding from NSERC, Canada). Research in A.S-P. group was supported by the European Research Council (ERC) under the European Union’s Horizon 2020 Research and Innovation Programme Grant Agreement (851647), the Spanish Ministry of Science and Innovation (PGC2018-098210-A-I00), the Centro de Excelencia Severo Ochoa scheme (SEV-2016-0571), and the Agencia Estatal de Investigacion. C.N. is supported by an FPI PhD fellowship from the Spanish Ministry of Economy, Industry and Competitiveness (MEIC). X.G-B. is supported by a Juan de la Cierva fellowship (FJC2018-036282-I) from the Spanish Ministry of Economy, Industry and Competitiveness (MEIC). I.R-T. was supported by a European Research Council Grant (616960). B.F.L. was supported by the Natural Sciences and Engineering Research Council of Canada (NSERC; RGPIN-2017-05411) and by the ‘Fonds de Recherche Nature et Technologie’, Quebec. P.L-G. and D.M. were supported by a Moore and Simons foundations grant (GBMF9739) and by European Research Council Advanced Grants (322669, 787904). Research in C.S. group was supported by the European Research Council (ERC) through project TACKLE (AdvGrant No. 695192).

## Author contributions

A.S.-P. conceived the project. X.G.-B., C.C., I.R.T., C.S., E.S. and A.S.-P. designed experiments and analytical strategies. C. N., T.P., M. A. and A.S.-P. performed experiments. X.G.-B., C.C., and A.S.-P analyzed the data. T.P., G.T., L.J.G., D.M., P.L-G. and B.F.L. provided biological samples/cultures and genomic data. All authors contributed to data interpretation. X.G.-B. and A.S.-P. wrote the manuscript with input from all authors.

## SUPPLEMENTARY FIGURES LEGENDS

**Supplementary Fig. 1. Histone classification and evolution. a,** Primary and secondary alignments of histone-fold containing proteins classified as canonical H2A, H2B, H3 and H4, based on identity to reference sequences in HistoneDB^48^. Pie plots represent the number of alignments to HistoneDB-annotated sequences, for the entire dataset (prokaryotic, eukaryotic and viral sequences, large pie plots in the inset) and the eukaryotic subset (smaller plots in the inset). For those proteins that align to more than one canonical histone or major variant (macroH2A, H2A.Z or cenH3), the scatter plots represent the relative identity between the primary (horizontal axis) and secondary alignment(s) (vertical axis). **b,** Aggregated counts of histone gene pairs, classified according to histone type and orientation. **c,** Presence of histone variants (left) and number of collinear pairs of histone-encoding genes (right) per species, classified according to their histone types and relative orientation (head-to-head, hh; head-to-tail, ht; and tail-to-tail, tt). Source data available in Supplementary Table 2. Histone variant classification is based on the highest-scoring HMM profile from HistoneDB. Asterisks colors in the macroH2A column indicate species where histone-less Macro domains orthologous to the macroH2A genes are found (see panel d). Lighter colors in the variant classification indicate ambiguously classified histones (i.e. cases in which the highest-scoring HMM profile exhibited a low bitscore, defined as a probability below 0.05 in the profile-wise distribution function of scaled bitscores; or cases in which the first-to-second ratio between high scoring profiles was below 1.01). **d,** Alignments of putatively conserved histone *N*-tails in archaea. Conserved aminoacids are color-coded according to chemical properties. Dots next to species names are color-coded according to taxonomy (same as **Fig. 2c**). **e,** Phylogenetic analysis of the Macro motif of macroH2A histones across eukaryotes, highlighting the macroH2A ortholog group (green), and, within this group, Macro-containing genes lacking histone domains (orange), and their protein domain architectures.

**Supplementary Fig. 2. Histone post-translational modifications. a,** Proteomics detection coverage (% of amino acids), number of hPTMs and number of hPTMs per covered position, for the best-covered histone in each species in our proteomics survey. **b,** Number of samples in which each histone-matching peptide with post-translational modifications (peptide spectral matches defined by *Proteome Discoverer*) has been identified, per species. For each species, we report the percentage of modified peptides found in more than one replicate. **c,** Number of samples in which histone-matching modified peptide has been identified, across all the samples from this study. The tree pie charts represent these distributions for all hPTMs, acetylations, and methylations. **d,** Evidence of hPTM conservation in the major histone variants H2A.Z and macroH2A (conserved positions only), as well as any position in the linker histones H1.

**Supplementary Fig. 3. Gene family counts. a-c,** Number of taxa within each lineage that contain chromatin-associated genes, for archaeal, bacterial (per phyla) or viral (per family) genomes. Numbers indicate the exact number of taxa. **d,** Number of genes encoding core domains that define chromatin-associated gene families per eukaryotic genome/transcriptome. Numbers indicate exact number of proteins.

**Supplementary Fig. 4. Evolutionary reconstruction and domain architecture conservation. a,** Species tree of eukaryotes used in the ancestral reconstruction analysis, with branch lengths calibrated to the gain/loss rates of Pfam domains (see Methods). Available in Supplementary Table 1. **b,** Conservation of archetypical protein domain architectures across orthogroups, in acetylases, deacetylases, methyltransferases, demethylases, remodellers and chaperones. In each heatmap, we indicate the fraction of genes within an orthogroup (rows) that contain a specific protein domain (columns). Domains in bold are catalytic (black) or reader (purple) functions. At the right of each heatmap, we summarize the presence/absence profile of each orthogroup across eukaryotic lineages (as listed in Fig. 1a).

**Supplementary Fig. 5. Evolution of the hPTM reader toolkit. a,** Pie plot representing the number of genes classified as part of the catalytic (acetylases, deacetylases, methyltransferases, demethylases, remodellers or chaperones) or reader families, or as both. The barplot at the right shows the most common reader domains in genes classified with both reader and catalytic functions**. b,** Pie plot representing the number of reader domain-encoding genes classified according to whether they contain one type of reader domain (e.g., PHD) or more than one (e.g., PHD + PWWP). The barplot at the right shows the most common combinations of reader domains among genes with multiple reader domains. **c,** Summary of gene family gains per reader family, with example cases highlighted in selected nodes. Node size is proportional to number of gains at 90% probability.

**Supplementary Fig. 6. Transposon-chromatin gene fusions. a,** Number of candidate fusion genes classified by the level of gene model validation evidence, based on contiguity of the gene model over the genome assembly (i.e. lack of poly-N stretches in the genomic region between the TE- and chromatin-associated domains), evidence of expression, and evidence of contiguous expression (see inset at the right). **b,** Summary of candidate gene fusions within each chromatin-associated gene family, divided by gene family. For each gene, we indicate their similarity to known TE families, presence of TE-associated domains, the evidence of gene model validity, and information on their gene structure (whether they are monoexonic or are located in clusters with other fusion genes). Source data available in Supplementary Table 6. **c,** Number of species with at least one valid fusion, divided by gene family. **d,** Mapping positions of RNA-seq reads supporting candidate gene-transposon fusions (selected examples from Fig. 5e). For each fusion, we show reads spanning the region along the spliced transcript that fully covers the transposon-associated domains (highlighted in green), the chromatin-associated domains, and the inter-domain region. Uninterrupted stretches of mapped positions between domains indicate the validity of a domain co-occurrence. For clarity purposes, reads mapping entirely within a single domain have been excluded from this visualization.

**Supplementary Fig. 7. Chromatin proteins in viruses. a-c,** Selected gene trees highlighting examples of eukaryotic- and prokaryotic-like viral homologs. **d,** Number of viral genes of each chromatin-associated gene family, classified according to their closest neighbours from cellular clades in gene tree analyses based on phylogenetic affinity scores (see Methods). Within each gene family, viral sequences are classified according to their PFAM domain architecture – the most common architecture being single-domain in most gene families except for remodellers and BIR readers. **e,***Id*., but classifying viral genes according to their phylogenetic affinity to eukaryotic orthology groups. Source data available in Supplementary Table 6.

**Supplementary Material 8. Phylogenetic analyses.** Collection of gene trees used to identify orthology groups for the eukaryotic chromatin toolkit. UFBS bootstrap supports rare indicated at each node. An annotated eukaryotic species tree is also included.

**Supplementary Material 9. Peptide sequences.** Collection of peptide sequences used to build gene trees of the eukaryotic chromatin toolkit.

## SUPPLEMENTARY TABLES LEGENDS

**Supplementary Table 1. Taxon sampling. a,** List of eukaryotic species used in the comparative genomic analyses, including species abbreviations, data sources for genome or transcriptome assemblies and annotations, and their taxonomic classification. **b,** List of gene expression datasets (SRA accession numbers) used for gene model validation analyses of candidate fusion genes. **c,** List of histone post-translational modification proteomics datasets used in this study (PRIDE accession numbers).

**Supplementary Table 2. Histone clusters and classification. a,** Pairs of collinear histone-encoding genes, including their genomic coordinates and relative orientation. **b,** List and sequences of archaeal HMfB histones with N-terminal tails (at least 10 aa before a complete globular domain). **c,** Classification of histone variants across eukaryotes.

**Supplementary Table 3. hPTM conservation. a-g,** Table of hPTMs identified in histones of the 26 eukaryotic species used in the comparative proteomics analysis, separated by histone type (canonical and major variants: H2A, H2B, H3, H4, macroH2A, H2A.Z, and H1). Each entry corresponds to a modified peptide, for which we specify modification coordinates along the peptide and relative to the consensus histone sequence (if available). We also indicate whether each peptide can be uniquely mapped to a conserved or non-conserved region in a canonical histone, or to specific histone variants. These tables also include entries for hPTMs reported in the literature (indicated as a cited source or as a specific UNIPROT entry; see Methods for a list of sources); in these cases, source peptides and associated data may not be available. **h,** hPTMs in Archaea.

**Supplementary Table 4. Gene family analysis. a,** List of gene classes analyzed in the comparative genomics analyses, including the PFAM protein domains used to retrieve homologs and search parameters. **b,** List of transposon-associated PFAM domains surveyed in the analyses of transposon-chromatin gene fusions.

**Supplementary Table 5. Evolution of the chromatin machinery in eukaryotes. a,** Summary of gene family evolutionary patterns in eukaryotes (*n* = 1,713 orthogroups). For each orthogroup, we indicate its gene and functional class, the number of members, species where it is present, and major eukaryotic lineages (Amoebozoa, Opisthokonta+Breviatea+Apusozoa, CRuMs, Ancyromonadida, Mala-wimonadidae, Archaeplastida+Cryptista, SAR+Haptista, Hemimastigophora, Discoba, and Metamonada), the probability of presence at the last eukaryotic common ancestor, the phylogenetic affinity of their closest homologs (other eukaryotic orthogroups, bacteria, archaea or viruses) and their average frequency amongst the 10 nearest neighbours of its member gene in phylogenetic trees (‘Phylogenetic affinity score’, see Methods); as well as its consensus protein domain architecture (present in at least 25% of its members). We also indicate the gene symbols of members from four model species: *H. sapiens, D. melanogaster*, *S. cerevisiae*, and *A. thaliana*. **b-c,** Probability of gain and loss of each gene family at extant and ancestral nodes along the eukaryotic phylogeny. **d,** Orthogroup assignments per gene.

**Supplementary Table 6. Transposon fusions and viral homology. a,** List of candidate fusions between chromatin-associated genes and transposons, including the phylogenetic classification of each gene (orthogroup), protein domain architectures, and the transcriptomics-level and gene model-level evidence supporting each fusion. **b,** List of chromatin-associated genes encoded by viral genomes, including their species of origin and a summary of their phylogenetic embedding among cellular species (specifically, which are its closest homologs in cellular genomes and the fraction of phylogenetic nearest neighbours they represent, the closest eukaryotic gene family among those close to eukaryotic genes in the gene trees, and the distance to the closest cellular homolog).

## Methods

### Eukaryotic cell culture and tissue sources

*Capsaspora owczarzaki* strain ATCC30864 filopodial cells were grown axenically in 5 ml flasks with ATCC medium 1034 (modified PYNFH medium) in an incubator at 23°C (Sebé-Pedrós et al., 2013a).

*Corallochytrium limacisporum* strain India was axenically grown in Difco Marine Broth medium at 23°C, *Creolimax fragrantissima* strain CH2 was axenically grown in Difco Marine Broth medium at 12°C, *Spizellomyces punctatus* strain DAOM BR117 was axenically grown in (0,5% yeast extract, 3% glycerol,1g/L K_2_HPO_4_, 0,5% EtOH) medium at 17°C, *Thecamonas trahens* strain ATCC50062 was grown in ATCC medium: 1525 Seawater 802 medium, *Chlamydomonas reinhardtii* strain CC-503 cw92 mt+ was axenically grown in Gibco TAP medium at 29°C, *Guillardia theta* strain CCMP2712 was axenically grown in L1+500uM NH_4_Cl medium at 18°C, *Emiliania huxleyi* strain CCMP1516 was grown in L1-Si medium at 18°C, *Thalassiosira pseudonana* strain CCMP1335 was axenically grown in L1 medium at 18°C, *Bigelowiella natans* strain CCMP2755 was axenically grown in L1-Si medium at 23°C, *Naegleria gruberi* strain ATCC30224 was axenically grown in ATCC medium 1034 (modified PYNFH medium) at 29°C, *Gefionella okellyi* strain 249 was grown in 15% Water Complete Cereal Grass Media (WC□CGM3) at 18°C and Fabomonas tropica strain NYK3C was grown in L1 + YT medium at 18°C. All cells were grown in 250 ml culture flasks.

In addition, we used frozen tissues/cells from the following species: *Homo sapiens* (ES cells, courtesy of Cecilia Ballaré, CRG), *Physcomitrella patens* (strain Gransden 2004, vegetative stage, courtesy of Josep Casacuberta, CRAG-CSIC), *Sycon ciliatum* (adult sponges sampled from Bergen, Norway, courtesy of Maja Adamska, ANU) and *Phytophthora infestans* (strain T30-4, courtesy of Harold J.G. Meijer, Wageningen University).

### Archaeal cell culture

Cultures of *Methanobrevibacter cuticularis* DSM 11139*, Methanospirillum stamsii* DSM 26304 and *Methanosarcina spelaei* DSM 26047 were purchased from the Deutsche Stammsammlung von Mikroorganismen und Zellkulturen GmbH (DSMZ), Braunschweig, Germany. Cultures were grown in closed batch in 50mL of defined media in 120mL serum bottles (La-Pha-Pack, Langerwehe, Germany). Growth was monitored as OD (600 nm; Analytik Jena, Specord 200 plus). *Methanobrevibacter cuticularis* was grown in modified *Methanobrevibacter cuticularis* medium DSMZ 734a (DSMZ 2014) omitting bovine rumen fluid, yeast extract and Na-resazurin at 1.5 bar overpressure H_2_CO_2_ (20 vol.-% CO2 in H_2_) at 37°C. As soon as a change in OD was observed, a constant agitation at 90rpm was applied. *Methanospirillum stamsii* was grown in modified *Methanobacterium* medium DSMZ 119 (DSMZ 2017) omitting sludge fluid, yeast extract and Naresazurin at 1 bar overpressure H_2_CO_2_ (20 vol.-% CO2 in H_2_) at 29°C, under constant agitation at 90rpm. *Methanosarcina spelaei* was grown in modified *Methanosarcina barkeri* medium DSMZ 120a (DSMZ 2014) omitting yeast extract and Na-resazurin at 1.5 bar overpressure H_2_CO_2_ (20 vol.-%CO2 in H_2_) at 33°C, under constant agitation at 90rpm. All gases were obtained from Air Liquide GmbH, Schwechat, Austria. *Nitrososphaera viennensis* EN76 was grown in continuous culture in a bioreactor as previously described^100^.

Cells were harvested via centrifugation at 21,000xg 4°C 1h (Thermo scientific, Sorvall Lynx 4000 centrifuge), the supernatant discarded and the resulting pellet resuspended in 1ml of spent medium, followed by another round of centrifugation at 21,000xg 4°C for 1h (Eppendorf, Centrifuge 5424R). Pellets were stored at −70°C. All archaeal histones were extracted as described below.

### Histone acid extraction

Starting material was a pellet of 50-100M cells (washed once with cold PBS) or a flash-frozen tissue homogenate in liquid nitrogen using a ceramic mortar grinder. Cells were washed first in 10ml of buffer I (10 mM TrisHCl pH 8, 10 mM MgCl_2_, 0.4M Sucrose). After 5min incubation, samples were centrifuged at 8.000g for 20min at 4°C and supernatant was removed. The resulting pellet was resuspended in 1.5ml of Buffer II (10 mM TrisHCl pH 8, 10 mM MgCl_2_, 0.25M Sucrose, 1% Triton X-100, 1% Igepal Ca-630) and incubated 15min on ice. In specific cases, cells at this stage were broken using a 2ml Dounce homogenizer (with Pestle B) or with a 20G syringe. Then samples were centrifuged at 15.000g for 10min at 4°C and supernatant was removed. The resulting pellet was then slowly resuspended in 300μL of Buffer III (10 mM TrisHCl pH 8, 2 mM MgCl_2_, 1.7M Sucrose, 1% Triton X-100) and then resulting resuspended nuclei were layered on top of another 300μL of Buffer III. Sample was centrifuged at 20.000g for 1h at 4°C and supernatant was removed, resulting in a nuclear pellet ready for acid histone extraction. All buffers were supplemented with spermidine (1:1000), beta-mercaptoethanol (1:1000), protease inhibitors (1x cOmplete cocktail Roche #11697498001, 1mM PMSF, 1:2000 Pepstatin), phosphatase inhibitors (1x phoSTOP cocktail Roche #4906845001) and deacetylase inhibitors (10mM Sodium butyrate).

For samples processed using a high-salt + HCl extraction protocol^101,102^, the pellet was resuspended in 500μL of High Salt Extraction Buffer (20 mM TrisHCl pH 7.4, CaCl_2_ 1M and protease, phosphatase and deacetylase inhibitors, same as above). Sample was incubated on ice for 30min and then pure HCl has added to a final 0.3N concentration (12.82μL to the initial 500μL). Samples were incubated for at least 2h on a rotor at 4°C and then centrifuged at 16.000g for 10min at 4°C to remove cellular/nuclear debris. The resulting supernatant containing solubilized histones was transferred to a clean 1.5ml tube and Trichloroacetic Acid (TCA) was added drop-wise to 25% final concentration (171μL TCA to an approximate initial 513μL sample) and left overnight at 4°C to precipitate histones. Samples were then centrifuged at 20.000g for 30min at 4°C and the supernatant removed. The pellet was then washed twice with 500μL of cold acetone and then dried for 20min at room temperature. Finally, clean histone pellets were resuspended in 30-50μL of ultrapure water. Protein concentration in the sample was measured using BCA and extraction was examined using an SDS-PAGE protein gel with Coomassie staining.

For samples processed using H_2_SO_4_^102^, the protocol was exactly the same except that 400μL 0.4N H_2_SO_4_ (freshly diluted) was used instead, with a similar incubation time of at least 2h at 4°C.

### Histone chemical derivatization

Histones samples were quantified by the BCA method and 10 μg of each sample were derivatized with propionic anhydride, digested with trypsin and derivatized again with phenylisocyanate as previously described^49^. Briefly, samples were dissolved in 9 μL of H2O and 1 μL of triethyl ammonium bicarbonate was added to bring the pH to 8.5. The propionic anhydride was prepared by adding 1 μL of propionic anhydride to 99 μL of H2O and 1 μL of propionic anhydride solution was added immediately to the samples with vortexing and incubation for 2 minutes. The reaction was quenched with 1 μL of 80mM hydroxylamine and samples were incubated at room temperature for 20 minutes. Tryptic digestion was performed for 3 h with 0.1 μg trypsin (Promega Sequencing Grade; Madison, WI) per sample. A 1% v/v solution of phenyl isocyanate (PIC) in acetonitrile was freshly prepared and 3 μl added to each sample (17 mM final concentration) and incubated for 60 min at 37 °C. Samples were acidified by adding 50 μL of 5% formic acid, vacuum dried and desalted with C18 ultramicrospin columns (The Nest Group, Inc, Southborough, MA).

### Liquid Chromatography-Tandem Mass Spectrometry Sample Acquisition

A 2-μg aliquot of the peptide mixture was analyzed using a LTQ-Orbitrap Fusion Lumos mass spectrometer (Thermo Fisher Scientific, San Jose, CA) coupled to an EASY-nLC 1000 (Thermo Fisher Scientific, San Jose, CA) with both collision induced dissociation (CID) and high energy collision dissociation (HCD) fragmentation.

Peptides were loaded directly onto the analytical column and were separated by reversed-phase chromatography using a 50-cm column with an inner diameter of 75 μm, packed with 2 μm C18 particles spectrometer (Thermo Scientific, San Jose, CA, USA) with a 90 min chromatographic gradient. The mass spectrometer was operated in positive ionization mode using a data dependent acquisition method. The “Top Speed” acquisition algorithm determined the number of selected precursor ions for fragmentation.

### Mass-spectrometry Data Analysis

Acquired data were analyzed using the Proteome Discoverer software suite (v2.0, Thermo Fisher Scientific), and the Mascot search engine (v2.6, Matrix Science^103^) was used for peptide identification using a double-search strategy. First, data were searched against each organism protein database plus the most common contaminants considering Propionylation on *N*-terminal, Propionylation on Lysines and Phenylisocyanate on *N*-terminal as variable modifications. Then a new database was generated with the proteins identified in the first search,, and a second search was done considering Propionylation on *N*-terminal, Propionylation on Lysines, Phenylisocyanate on *N*-terminal, Dimethyl lysine, trimethyl lysine, propionyl + methyl lysine, acetyl lysine, crotonyl lysine as variable modifications. Precursor ion mass tolerance of 7 ppm at the MS1 level was used, and up to 5 missed cleavages for trypsin were allowed. False discovery rate (FDR) in peptide identification was set to a maximum of 5%. The identified peptides were filtered by mascot ion score higher than 20 and only PTMs with a localization score ptmRS^104^ higher than 45 were considered. The raw proteomics data have been deposited to the PRIDE^105^ repository with the dataset identifier PXD031991.

### Analysis of hPTM conservation

#### Identification of canonical and variant histones

We classified histone protein domains from a database of eukaryotic, prokaryotic and viral sequences (see details below) according to their similarity to known canonical (H2A, H2B, H3, H4) and variant histones (e.g., H2A.Z, macroH2A, cenH3 or H3.3), as well as other gene families with histone-like protein folds (e.g., the transcription factors DR1, DRAP1, NFYB/C, POLE3/4, SOS, TAF, or CHRAC). To that end, we used *diamond* to perform local alignments of each histone domain against (*i*) a set of curated histone variants obtained from HistoneDB 2.0^48^, and (*ii*) annotated each domain according to the best hit in the reference database, which allowed us to classify histone fold-containing proteins as canonical histones (H2A, H2B, H3, H4) or their main variants (H2A.Z, macroH2A and cenH3). This best-hit strategy performs well in distinguishing canonical histones from each other, as well as each canonical histone from its main variants (H3 from cenH3, and H2A from H2A.Z and macroH2A; **Supplementary Fig. 1a**).

Then, we built a graph of pairwise similarity between histones, with edges weighted by the alignment bitscore (discarding edges with bitscore < 20). We created visualisations of each connected component in this graph using the spring layout algorithm implemented in the *networkx* 2.4 Python library (100 iterations, weighted by alignment bitscore)^106^. We selected the four connected components in the graph that matched the four canonical eukaryotic histones (H2A, H2B, H3, H4; discarding edges with bitscore < 20), retrieved the protein sequences for each of them, aligned them using *mafft* (E-INS-i mode, 1,000 iterations)^107^, and built phylogenetic trees with *IQ-TREE* 2.1.0 (*-fast* mode)^108^.

#### Identification of hPTM homology

We retrieved the protein sequences of the canonical histones identified in each of the 26 species and we used them for the proteomic analysis of hPTMs, and aligned them using *mafft* (*G-INS-i* mode, up to 10,000 refinement iterations). For this subset of species, histone class identity was cross-referenced with the HistoneDB search tool. Then, we manually aligned the peptides mapping onto these proteins to identify the position of each hPTM along a consensus alignment. In the case of H3, H4, and macroH2A, the majority of alignment positions were conserved across most eukaryotes in our dataset, and we used a consensus numbering scheme. In the case of H2A, H2A.Z, and H2B, non-conserved insertions and deletions at the N-terminal tail precluded the use of a paneukaryotic numbering scheme. Instead, we reported hPTM positions based on the human homolog (if possible), or relative to taxonomically restricted conserved positions. In cases where position-wise homology could not be established, we grouped multiple aminoacids into stretches of unclear homology, which we report separately from conserved positions (question mark symbols in **Fig. 1**). The complete list of hPTMs and their position-wise coordinates relative to the consensus alignment is available in **Supplementary Table 3**.

Furthermore, we also reported the presence (in any position) of modifications in less-conserved histone variants, as well as the linker histone H1.

In addition to the 19 used in our proteomics survey, we also included previously published hPTM data from the following species (**Supplementary Table 1c**): the brown alga *Ectocarpus siliculosus*^43^, the diatom *Phaeodactylum tricornutum*^109^, the ciliate *Tetrahymena thermophila*^46,110–112^, the ascomycete *Neurospora crassa*^113^. *Saccharomyces cerevisiae* and *Schizosaccharomyces pombe*^46^, and the plant *Arabidopsis thaliana*^114–116^. When available in public repositories, we reanalysed these datasets using the strategy described above. Finally, we also complemented our own proteomics data using previously published hPTM data from *Homo sapiens*^46,117–120^ and *Capsaspora owczarzaki*^42^.

### Comparative genomics analysis of chromatin-associated proteins

#### Data retrieval

We identified homologs of gene families associated with eukaryotic chromatin, using a database of predicted proteomes from a selection of eukaryotic species from all major super-groups (*n* = 172 species; see **Supplementary Table 1** for their taxonomic classification and data sources), as well as archaeal and viral peptides available in the NCBI *nr* peptide collection (as of 25th of April, 2020) and bacterial peptides available in RefSeq (release 99, 11th May, 2020). The database of viral sequences was complemented with peptides from 501 genomes of nucleocytoplasmic large DNA viruses^121^.

#### Gene family searches

We defined 61 gene classes associated with eukaryotic chromatin, based on HMM models obtained from the Pfam database (release 33.0)^122^. This list included canonical and linker histones (*n* = 2 families), chromatin-specific lysine acetylases (*n* = 5), deacetylases (*n* = 2), methyltransferases (*n* = 2), demethylases (*n* = 2), chromatin readers (*n* = 16), remodellers (*n* = 1) and chaperones (*n* = 13), as well as multiple families associated with the Polycomb complexes (*n* = 18). The complete list of gene families, including the associated HMM models, is available in **Supplementary Table 4**.

For each gene family, we retrieved all homologs from the eukaryotic, archaeal, bacterial and viral databases using the *hmmsearch* tool from the *HMMER* 3.3 toolkit^123^ and the gathering threshold defined in each Pfam HMM model. We recorded the taxonomic profile of each homolog.

#### Orthology identification

We aimed to identify groups of orthologs within each of the 61 chromatin-associated gene families using targeted phylogenetic analyses. We followed the following strategy for each of the 59 sets of eukaryotic genes. First, we partitioned each set into one or more homology groups based on pairwise local sequence alignments using *diamond* 0.9.36.137 (high sensitivity all-to-all search)^124^, followed by clustering of the resulting pairwise alignments graph with *MCL* 14.137 (*--abc* mode)^125^, using low inflation values (see **Supplementary Table 4**) to favour inclusive groupings. Second, we performed multiple sequence alignments of each homology group with *mafft* 7.471^107^ under the E-INS-i mode (optimised for multiple conserved regions), running up to 10,000 refinement iterations. Third, we trimmed the resulting multiple sequence alignments using *clip-kit* 0.1 (*kpic-gappy* mode)^126^. Fourth, we built phylogenetic trees for each trimmed alignment using *IQ-TREE* 2.1.0^108^, selecting the best-fitting evolutionary model using its *ModelTest* module (according to the Bayesian Information Criterion) and using 1,000 UFBS bootstrap supports ^127^. Each tree was run for up to 10,000 iterations until convergence was attained (at the 0.999 correlation coefficient threshold, and for at least 200 iterations).

Then, we parsed the species composition of each gene tree in order to identify groups of orthologous proteins using the *POSSVM* pipeline^128^. Specifically, we used the species overlap algorithm^129^ implemented in the *ETE* toolkit 3.1.1^130^, which identifies pairs of orthologous genes in a phylogenetic tree by examining the species composition of each subtree, and classifying internal nodes as paralogy nodes (if there is overlap in the species composition between each of its two descendant subtrees) or orthology nodes (if there is no overlap). Pairs of genes linked by an orthology node are then recorded as orthology pairs. In our analysis, we used an overlap threshold=0 (i.e. any species composition overlap between the two descendant subtrees is classified as a paralogy event). The resulting list of pairwise orthology relationships between genes was clustered into groups of orthologs (orthogroups) using *MCL*. We further annotated each orthogroup with a string denoting the gene symbols of the human proteins therein (if any).

Overall, we classified 51,426 proteins from 61 gene classes (defined by protein structural domains), divided into 242 gene trees and 1,713 gene families (orthogroups). The source peptide sequences and gene trees used for these analyses are available in **Supplementary Material 7 and 8**.

#### Ancestral reconstruction of gene content

We inferred the presence, gain and loss of each orthogroup along the eukaryotic tree of life, using a phylogenetic birth-and-death model^131^ implemented in *Count*^132^. This tool takes a numeric profile of gene family presence/absence in extant species (172 in our dataset) and a phylogenetic tree defining their evolutionary relationships, and infers the probabilities of gain and loss of each family at each ancestral node along the tree.

First we trained the probabilistic model in *Count*. As a training set, we used a random sample of 1,000 PFAM domains annotated in the 172 species of interest (restricting the sampling to domains present in at least 5% of species). The final model consists of gain, loss and transfer rates with two Γ categories each, and a constant duplication rate (given that we only recorded gene presence/absence, duplication events are not included in our downstream analyses). This model was obtained in three sequential rounds of training, so as to sequentially add zero, one and two Γ categories to each evolutionary rate. Each round consisted of up to 100 iterations, and stopped when the relative change in the model log-likelihood fell by 0.1% in two consecutive rounds. The final evolutionary rates and the Newick-formatted species tree used in this step are available in the **Supplementary Table 1 and Supplementary Fig. 3a**.

Second, we calculated the posterior probability of gain, loss and presence of each orthogroup in our dataset with *Count*. The aggregated counts of gains and losses of the various classes of chromatin-associated proteins (acetylases, deacetylases, methyltransferases, demethylases, readers and remodellers) along the eukaryotic tree were obtained by summing the probabilities of gain, presence or loss of all orthogroups of a given class at each ancestral node. To investigate the evolutionary histories of specific orthogroups at a given node in the tree, we applied a probability threshold of 0.9 (for presence) or 0.5 (to identify the most probable gain and loss node). The *Count* model was not able to calculate ancestral probabilities for a few orthogroups with widespread phylogenetic distributions, due to violations of the birth-and-death model (25 out of 1,713 families). In order to be able to report presence probabilities in the LECA for these orthogroups, we inferred their presence in this ancestor using the Wagner parsimony procedure implemented in *Count* with a gain-to-loss penalty *g* = 5, and recorded their presence as binary values (0/1) accordingly.

#### Protein domain architecture analyses

We annotated the Pfam domains present in each protein from the gene classes listed in **Supplementary Table 4**, using *Pfamscan* 1.6-3 and the Pfam 33.0 database^122^. We visualized the networks of protein domain co-occurrence from the point of view of the core domain(s) that define each gene class, using the *networkx* Python library (version 2.4)^106^. Specifically, we built a graph where each node represented ‘accessory’ domains (i.e. domains that co-occur with the ‘core’ domain that defines given gene class), node size reflected number of co-occurrences with the ‘core’ domain, and edges reflected co-occurrences between accessory domains. We identified communities of frequently co-occurring accessory domains using the label propagation algorithm implemented in *networkx (communities* submodule), which we used as a basis to manually annotate groups of co-occurring domains of interest (**Fig. 5C**). Network visualizations were created using the *NEATO* spring layout algorithm from the *Graphviz* 2.40.1 Python library^133^.

In parallel, we also recorded the presence of Pfam domains within individual orthogroups, and their taxonomic distribution.

#### Prokaryotic roots of the eukaryotic chromatin machinery

We retrieved all eukaryotic domains from gene class shared with prokaryotes (Histones, Acetyltransf_1, GNAT_acetyltr_2, MOZ_SAS, Hist_deacetyl, SIR2, DOT1, SET, CupinJmjC, ING, MBT, PWWP and SNF2_N), collapsing identical sequences at 100% similarity with *CD-HIT* 4.8.1^134^, and identified their closest homologs amongst the corresponding archaea and bacteria protein domain sets, using *diamond* local alignments (high sensitivity search). The archaeal and bacterial protein sets were also reduced with *CD-HIT* (at 95% and 90% sequence similarity, respectively). Each set of sequences was then partitioned into low-granularity homology clusters using the *MCL*-based strategy described above (inflation *I* = 1.2), and a phylogenetic tree was then constructed from each homology cluster with *IQ-TREE* (as described above).

Then, we mapped each eukaryotic gene to its orthogroup (obtained from eukaryotic-only analyses, see above) and used the distribution of phylogenetic distances from the prokaryotic+eukaryotic gene trees to classify them according to their similarity to (*i*) eukaryotic genes in other orthogroups, (*ii*) archaeal homologs, or (*iii*) bacterial homologs. Specifically, we used a majority-voting procedure in which we recorded the number of sequences of eukaryotic, archaeal or bacterial origin amongst the ten nearest neighbors of each gene (measuring intergenic distances as substitutions per site), and assigned the most common taxonomic group as the ‘closest’ homolog of that gene (minimum 50% agreement). This fraction is termed ‘Phylogenetic affinity score’ and reported in **Supplementary Table 5**. The pairwise distances were obtained from each gene tree using the cophenetic distance method in the *cophenetic.phylo* utility of the *ape* 5.4 *R* library^135^.

#### Characterisation of fusions with transposon-associated domains

We retrieved all classified genes from our eukaryotic dataset that contained transposon-associated Pfam domains (version 33.0), using a list compiled from^68,136^ (complete list in **Supplementary Table 4**), totaling 823 candidate fusions from 91 species (listed in **Supplementary Table 6**). We annotated these genes to their most similar known TE element by aligning them against the Dfam 3.3 database^137^ using the *tblastn* program in *BLAST* 2.2.31^138^.

We validated each candidate fusion using the following criteria: (*i*) contiguity of the gene model on the genome assembly, i.e., recording which genes were interrupted by poly-*N* stretches (which might indicate an incorrect gene model); (*ii*) evidence of expression in at least one sample from a range of publicly available transcriptomic experiments (from the NCBI SRA repository); (*iii*) evidence of contiguous expression, i.e., whether an expressed transcript had mapped reads along the entire region located between the ‘core’ and ‘TE-associated’ domains; (iv) we also recorded the number of exons per gene; and (*v*) located near any other candidate fusion gene in the genome.

The list of SRA experiments used for these validation steps is available in **Supplementary Table 1**. This list includes 64 out of 91 species for which transcriptomics datasets are publicly available, and covers 768 out of the 822 TE fusion candidates (93%). RNA-seq read mapping was performed with *bwa mem* 0.7.17-r1188^139^ using the complete set of spliced transcripts of each species as the reference database. We used *bedtools* 2.29.2^140^ to identify poly-*N* stretches in the genome assembly (assembly contiguity criterion). We identified regions of low coverage along the transcript sequence (expression contiguity criterion) using the *bedtools genomecov* utility, requiring that the coverage along both domains involved in each fusion and their intermediate regions be higher or equal to two reads.

#### Analysis of viral homologs

We investigated the homology of the viral chromatin-associated genes (which included 19 out of 61 families present in our survey) using joint phylogenetic analyses of protein domains from virus, prokaryotic and eukaryotic genes. We used the same method described above to investigate the prokaryotic roots of eukaryotic gene classes: we aligned viral domains against a database of cellular homologs (high sensitivity *diamond* search), followed by low-granularity *MCL* clustering (inflation *I* = 1.2) and phylogenetic tree building (*IQ-TREE*). Then, we used the same majority-voting procedure described above to classify viral homologs according to their similarity to eukaryotic, archaeal or bacterial gene families based on their distribution of phylogenetic distances. For viral genes that were most similar to eukaryotic genes, we used the same procedure to map them to their closest eukaryotic orthogroup.

The complete list of viral genes and their phylogenetic annotation is available in **Supplementary Table 6**. Out of 2,163 viral genes in our dataset, 2,144 could be annotated as similar to a particular cellular group using this procedure (99.1%), and the majority of these genes had a high agreement in the annotations of their nearest neighbors (2,096 with ≥50% agreement; 1,449 with ≥90% agreement).

In the case of viral histones, we built a separate phylogeny with a few modifications in our protocol: (*i*) we used additional viral genes obtained from^71^ as a reference; (*ii*) we omitted the *CD-HIT* reduction and *MCL* partitioning steps, and jointly analyzed the entire set of homologs instead; and (*iii*) in the phylogenetic reconstruction step, we used the approximate Bayes posterior probabilities^141^ implemented in *IQ-TREE*.

#### Identification of archaeal N-terminal histone tails

We retrieved all archaeal histone domains classified belonging to the HMfB-like connected component in **Fig. 1b**, and retained those that fulfilled the following criteria: (i) contained a complete CBFD_NFYB_HMF domain according to the *hmmscan* search (defined as an alignment starting at least at the 10th position of the HMM model, and up to the 55th position; the HMM model contains 65 positions); and (ii) the predicted tail (*N*-terminal to the core domain boundaries defined by *hmmscan*) was at least 10 residues long. 84 genes passed these filters, including three *N*-terminal containing histones previously identified by Henneman *et al.^55^*. A complete list is available in **Supplementary Table 2**. We manually examined the sequences of archaeal tails and aligned four sets of similar histones with *mafft G-INS-i* (**Supplementary Fig. 1d**). Alignments were plotted using the *msa* 1.24.0 library in *R*^142^.

### Data and Code Availability

The mass spectrometry proteomics data have been deposited to the ProteomeXchange Consortium via the PRIDE partner repository with the dataset identifier PXD031991. Code for reproducing the analysis is available in our lab Github repository (https://github.com/sebepedroslab/chromatin-evolution-analysis).

## References

1. Struhl, K. Fundamentally different logic of gene regulation in eukaryotes and prokaryotes. Cell 98, 1–4 (1999).

2. Kornberg, R. D. & Lorch, Y. Primary Role of the Nucleosome. Mol. Cell 79, 371–375 (2020).

3. Jenuwein, T. & Allis, C. D. Translating the Histone Code. Science 293, 1074–1080 (2001).

4. Berger, S. L. The complex language of chromatin regulation during transcription. Nature 447, 407–12 (2007).

5. Banaszynski, L. a, Allis, C. D. & Lewis, P. W. Histone variants in metazoan development. Dev. Cell 19, 662–74 (2010).

6. Allis, C. D. & Jenuwein, T. The molecular hallmarks of epigenetic control. Nat. Rev. Genet. 1, (2016).

7. Sultana, T. et al. The Landscape of L1 Retrotransposons in the Human Genome Is Shaped by Pre-insertion Sequence Biases and Post-insertion Selection. Mol. Cell 74, 555–570.e7 (2019).

8. Gangadharan, S., Mularoni, L., Fain-Thornton, J., Wheelan, S. J. & Craig, N. L. DNA transposon Hermes inserts into DNA in nucleosome-free regions in vivo. Proc. Natl. Acad. Sci. 107, 21966–21972 (2010).

9. Shinn, P. et al. HIV-1 Integration in the Human Genome Favors Active Genes and Local Hotspots. Cell 110, 521–529 (2002).

10. Goodier, J. L. Restricting retrotransposons: a review. Mob. DNA 7, 16 (2016).

11. Molaro, A. & Malik, H. S. Hide and seek: how chromatin-based pathways silence retroelements in the mammalian germline. Curr. Opin. Genet. Dev. 37, 51–58 (2016).

12. Malik, H. S. & Henikoff, S. Phylogenomics of the nucleosome. Nat. Struct. Biol. 10, 882–91 (2003).

13. Talbert, P. B. & Henikoff, S. Histone variants--ancient wrap artists of the epigenome. Nat. Rev. Mol. Cell Biol. 11, 264–75 (2010).

14. Soboleva, T. a., Nekrasov, M., Ryan, D. P. & Tremethick, D. J. Histone variants at the transcription start-site. Trends Genet. 30, 199–209 (2014).

15. Zink, L.-M. & Hake, S. B. Histone variants: nuclear function and disease. Curr. Opin. Genet. Dev. 37, 82–89 (2016).

16. Weber, C. M. & Henikoff, S. Histone variants: dynamic punctuation in transcription. Genes Dev. 28, 672–82 (2014).

17. Borg, M., Jiang, D. & Berger, F. Histone variants take center stage in shaping the epigenome. Curr. Opin. Plant Biol. 61, 101991 (2021).

18. Zentner, G. E. & Henikoff, S. Regulation of nucleosome dynamics by histone modifications. Nat. Struct. Mol. Biol. 20, 259–66 (2013).

19. Campos, E. I. & Reinberg, D. Histones: annotating chromatin. Annu. Rev. Genet. 43, 559–99 (2009).

20. Strahl, B. D. & Allis, C. D. The language of covalent histone modifications. Nature 403, 41–45 (2000).

21. Bannister, A. J. & Kouzarides, T. Regulation of chromatin by histone modifications. Cell Res. 21, 381–95 (2011).

22. Talbert, P. B. & Henikoff, S. The Yin and Yang of Histone Marks in Transcription. Annu. Rev. Genomics Hum. Genet. 22, 147–170 (2021).

23. Taverna, S. D., Li, H., Ruthenburg, A. J., Allis, C. D. & Patel, D. J. How chromatin-binding modules interpret histone modifications: lessons from professional pocket pickers. Nat. Struct. Mol. Biol. 14, 1025–40 (2007).

24. Musselman, C. a, Lalonde, M.-E., Côté, J. & Kutateladze, T. G. Perceiving the epigenetic landscape through histone readers. Nat. Struct. Mol. Biol. 19, 1218–27 (2012).

25. Gurard-Levin, Z. a, Quivy, J.-P. & Almouzni, G. Histone Chaperones: Assisting Histone Traffic and Nucleosome Dynamics. Annu. Rev. Biochem. 83, 487–517 (2014).

26. Burgess, R. J. & Zhang, Z. Histone chaperones in nucleosome assembly and human disease. Nat. Struct. Mol. Biol. 20, 14–22 (2013).

27. Koster, M. J. E., Snel, B. & Timmers, H. T. M. Genesis of Chromatin and Transcription Dynamics in the Origin of Species. Cell 161, 724–736 (2015).

28. Hargreaves, D. C. & Crabtree, G. R. ATP-dependent chromatin remodeling: genetics, genomics and mechanisms. Cell Res. 21, 396–420 (2011).

29. Gornik, S. G. et al. Loss of nucleosomal DNA condensation coincides with appearance of a novel nuclear protein in dinoflagellates. Curr. Biol. 22, 2303–12 (2012).

30. Mattiroli, F. et al. Structure of histone-based chromatin in Archaea. Science 357, 609–612 (2017).

31. Warnecke, T., Becker, E. a, Facciotti, M. T., Nislow, C. & Lehner, B. Conserved substitution patterns around nucleosome footprints in eukaryotes and Archaea derive from frequent nucleosome repositioning through evolution. PLoS Comput. Biol. 9, e1003373 (2013).

32. Ammar, R. et al. Chromatin is an ancient innovation conserved between Archaea and Eukarya. Elife 1, e00078 (2012).

33. Rojec, M., Hocher, A., Merkenschlager, M. & Warnecke, T. Chromatinization of Escherichia coli with archaeal histones. bioRxiv 660035 (2019) doi:10.1101/660035.

34. Forbes, A. J. et al. Targeted analysis and discovery of posttranslational modifications in proteins from methanogenic archaea by top-down MS. Proc. Natl. Acad. Sci. U. S. A. 101, 2678–83 (2004).

35. Weidenbach, K. et al. Deletion of the archaeal histone in Methanosarcina mazei Gö1 results in reduced growth and genomic transcription. Mol. Microbiol. 67, 662–671 (2008).

36. Talbert, P. B., Meers, M. P. & Henikoff, S. Old cogs, new tricks: the evolution of gene expression in a chromatin context. Nat. Rev. Genet. (2019) doi:10.1038/s41576-019-0105-7.

37. de Mendoza, A. & Sebe-Pedros, A. Origin and evolution of eukaryotic transcription factors. Curr. Opin. Genet. Dev. 59, 25–32 (2019).

38. Schwaiger, M. et al. Evolutionary conservation of the eumetazoan gene regulatory landscape. Genome Res. 24, 639–650 (2014).

39. Sebé-Pedrós, A. et al. Early metazoan cell type diversity and the evolution of multicellular gene regulation. Nat. Ecol. Evol. 2, 1176–1188 (2018).

40. Connolly, L. R., Smith, K. M. & Freitag, M. The Fusarium graminearum Histone H3 K27 Methyltransferase KMT6 Regulates Development and Expression of Secondary Metabolite Gene Clusters. PLoS Genet. 9, e1003916 (2013).

41. Jamieson, K., Rountree, M. R., Lewis, Z. a, Stajich, J. E. & Selker, E. U. Regional control of histone H3 lysine 27 methylation in Neurospora. Proc. Natl. Acad. Sci. 110, 6027–6032 (2013).

42. Sebé-Pedrós, A. et al. The Dynamic Regulatory Genome of Capsaspora and the Origin of Animal Multicellularity. Cell 165, 1224–1237 (2016).

43. Bourdareau, S. et al. Histone modifications during the life cycle of the brown alga Ectocarpus. Genome Biol. 22, 12 (2021).

44. Wang, S. Y. et al. Role of epigenetics in unicellular to multicellular transition in Dictyostelium. Genome Biol. 22, 134 (2021).

45. Taverna, S. D., Coyne, R. S. & Allis, C. D. Methylation of Histone H3 at Lysine 9 Targets Programmed DNA Elimination in Tetrahymena University of Virginia Health System. Cell 110, 701–711 (2002).

46. Garcia, B. a et al. Organismal differences in post-translational modifications in histones H3 and H4. J. Biol. Chem. 282, 7641–55 (2007).

47. Drinnenberg, I. A. et al. EvoChromo: towards a synthesis of chromatin biology and evolution. Development 146, dev178962 (2019).

48. Draizen, E. J. et al. HistoneDB 2.0: a histone database with variants—an integrated resource to explore histones and their variants. Database 2016, baw014 (2016).

49. Maile, T. M. et al. Mass Spectrometric Quantification of Histone Post-translational Modifications by a Hybrid Chemical Labeling Method. Mol. Cell. Proteomics 14, 1148–1158 (2015).

50. Li, B., Carey, M. & Workman, J. L. The Role of Chromatin during Transcription. Cell 128, 707–719 (2007).

51. Rajagopal, N. et al. Distinct and Predictive Histone Lysine Acetylation Patterns at Promoters, Enhancers, and Gene Bodies. G3 Genes/Genomes/Genetics 4, 2051–2063 (2014).

52. Koonin, E. V. & Yutin, N. The Dispersed Archaeal Eukaryome and the Complex Archaeal Ancestor of Eukaryotes. Cold Spring Harb. Perspect. Biol. 6, a016188–a016188 (2014).

53. Sandman, K. & Reeve, J. N. Archaeal histones and the origin of the histone fold. Curr. Opin. Microbiol. 9, 520–5 (2006).

54. Pereira, S. L., Grayling, R. a, Lurz, R. & Reeve, J. N. Archaeal nucleosomes. Proc. Natl. Acad. Sci. U. S. A. 94, 12633–7 (1997).

55. Henneman, B., van Emmerik, C., van Ingen, H. & Dame, R. T. Structure and function of archaeal histones. PLOS Genet. 14, e1007582 (2018).

56. Imachi, H. et al. Isolation of an archaeon at the prokaryote-eukaryote interface. bioRxiv 726976 (2019) doi:10.1101/726976.

57. Spang, A. et al. Complex archaea that bridge the gap between prokaryotes and eukaryotes. Nature 521, 173–179 (2015).

58. Zaremba-Niedzwiedzka, K. et al. Asgard archaea illuminate the origin of eukaryotic cellular complexity. Nature 541, 353–358 (2017).

59. Da Cunha, V., Gaia, M., Nasir, A. & Forterre, P. Asgard archaea do not close the debate about the universal tree of life topology. PLOS Genet. 14, e1007215 (2018).

60. Alva, V. & Lupas, A. N. Histones predate the split between bacteria and archaea. Bioinformatics 35, 2349–2353 (2019).

61. Allis, C. D. et al. New Nomenclature for Chromatin-Modifying Enzymes. Cell 131, 633–636 (2007).

62. Wu, F. et al. Unique mobile elements and scalable gene flow at the prokaryote–eukaryote boundary revealed by circularized Asgard archaea genomes. Nat. Microbiol. 7, 200–212 (2022).

63. Schuettengruber, B., Bourbon, H.-M., Di Croce, L. & Cavalli, G. Genome Regulation by Polycomb and Trithorax: 70 Years and Counting. Cell 171, 34–57 (2017).

64. Dion, M. F., Altschuler, S. J., Wu, L. F. & Rando, O. J. Genomic characterization reveals a simple histone H4 acetylation code. Proc. Natl. Acad. Sci. 102, 5501–5506 (2005).

65. de Jong, J. et al. Chromatin Landscapes of Retroviral and Transposon Integration Profiles. PLoS Genet. 10, e1004250 (2014).

66. Sultana, T., Zamborlini, A., Cristofari, G. & Lesage, P. Integration site selection by retroviruses and transposable elements in eukaryotes. Nat. Rev. Genet. 18, 292–308 (2017).

67. Gao, X., Hou, Y., Ebina, H., Levin, H. L. & Voytas, D. F. Chromodomains direct integration of retrotransposons to heterochromatin. Genome Res. 18, 359–369 (2008).

68. Cosby, R. L. et al. Recurrent evolution of vertebrate transcription factors by transposase capture. Science 371, eabc6405 (2021).

69. Cordaux, R., Udit, S., Batzer, M. A. & Feschotte, C. Birth of a chimeric primate gene by capture of the transposase gene from a mobile element. Proc. Natl. Acad. Sci. 103, 8101 LP–8106 (2006).

70. Fiedler, M. et al. Decoding of Methylated Histone H3 Tail by the Pygo-BCL9 Wnt Signaling Complex. Mol. Cell 30, 507–518 (2008).

71. Erives, A. J. Phylogenetic analysis of the core histone doublet and DNA topo II genes of Marseilleviridae: evidence of proto-eukaryotic provenance. Epigenetics Chromatin 10, 55 (2017).

72. Liu, Y. et al. Virus-encoded histone doublets are essential and form nucleosome-like structures. Cell 184, 4237–4250.e19 (2021).

73. Valencia-Sánchez, M. I. et al. The structure of a virus-encoded nucleosome. Nat. Struct. Mol. Biol. 28, 413–417 (2021).

74. Iyer, L. M., Balaji, S., Koonin, E. V & Aravind, L. Evolutionary genomics of nucleo-cytoplasmic large DNA viruses. Virus Res. 117, 156–184 (2006).

75. Nagamine, T. Apoptotic arms races in insect baculovirus coevolution. Physiol. Entomol. phen.12371 (2021) doi:10.1111/phen.12371.

76. Starrett, G. J. et al. Adintoviruses: An Animal-Tropic Family of Midsize Eukaryotic Linear dsDNA (MELD) Viruses. bioRxiv 697771 (2020) doi:10.1101/697771.

77. Hocher, A. et al. Growth temperature is the principal driver of chromatinization in archaea. bioRxiv 2021.07.08.451601 (2021) doi:10.1101/2021.07.08.451601.

78. Alpha-Bazin, B. et al. Lysine-specific acetylated proteome from the archaeon Thermococcus gammatolerans reveals the presence of acetylated histones. J. Proteomics 232, 104044 (2021).

79. Eme, L., Spang, A., Lombard, J., Stairs, C. W. & Ettema, T. J. G. Archaea and the origin of eukaryotes. Nat. Rev. Microbiol. 15, 711–723 (2017).

80. Akil, C. & Robinson, R. C. Genomes of Asgard archaea encode profilins that regulate actin. Nature 562, 439–443 (2018).

81. Koonin, E. V The origin and early evolution of eukaryotes in the light of phylogenomics. Genome Biol. 11, 209 (2010).

82. Sebé-Pedrós, A., Grau-Bové, X., Richards, T. a & Ruiz-Trillo, I. Evolution and Classification of Myosins, a Paneukaryotic Whole-Genome Approach. Genome Biol. Evol. 6, 290–305 (2014).

83. Richards, T. A. & Cavalier-Smith, T. Myosin domain evolution and the primary divergence of eukaryotes. Nature 436, 1113–8 (2005).

84. Wickstead, B., Gull, K. & Richards, T. Patterns of kinesin evolution reveal a complex ancestral eukaryote with a multifunctional cytoskeleton. BMC Evol. Biol. 10, 110 (2010).

85. Dacks, J. B. & Field, M. C. Evolution of the eukaryotic membrane-trafficking system: origin, tempo and mode. J. Cell Sci. 120, 2977–85 (2007).

86. Collins, L. & Penny, D. Complex Spliceosomal Organization Ancestral to Extant Eukaryotes. Mol. Biol. Evol. 22, 1053–1066 (2005).

87. Grau-Bové, X., Sebé-Pedrós, A. & Ruiz-Trillo, I. The Eukaryotic Ancestor Had a Complex Ubiquitin Signaling System of Archaeal Origin. Mol. Biol. Evol. 32, 726–739 (2015).

88. Kundaje, A. et al. Integrative analysis of 111 reference human epigenomes. Nature 518, 317–330 (2015).

89. Ho, J. W. K. et al. Comparative analysis of metazoan chromatin organization. Nature 512, 449–452 (2014).

90. Montgomery, S. A. et al. Chromatin Organization in Early Land Plants Reveals an Ancestral Association between H3K27me3, Transposons, and Constitutive Heterochromatin. Curr. Biol. 30, 573–588.e7 (2020).

91. Frapporti, A. et al. The Polycomb protein Ezl1 mediates H3K9 and H3K27 methylation to repress transposable elements in Paramecium. Nat. Commun. 10, 2710 (2019).

92. Lennartsson, A. & Ekwall, K. Histone modification patterns and epigenetic codes. Biochim. Biophys. Acta 1790, 863–8 (2009).

93. Peterson, C. L. & Laniel, M.-A. Histones and histone modifications. Curr. Biol. 14, R546–51 (2004).

94. Rando, O. J. Combinatorial complexity in chromatin structure and function: revisiting the histone code. Curr. Opin. Genet. Dev. 22, 148–155 (2012).

95. de Mendoza, A., Pflueger, J. & Lister, R. Capture of a functionally active methyl-CpG binding domain by an arthropod retrotransposon family. Genome Res. 29, 1277–1286 (2019).

96. De Mendoza, A. et al. Recurrent acquisition of cytosine methyltransferases into eukaryotic retrotransposons. Nat. Commun. 9, 1–11 (2018).

97. Ji, X. et al. Chromatin proteomic profiling reveals novel proteins associated with histone-marked genomic regions. Proc. Natl. Acad. Sci. U. S. A. 112, 3841–3846 (2015).

98. Wierer, M. & Mann, M. Proteomics to study DNA-bound and chromatin-associated gene regulatory complexes. Hum. Mol. Genet. 25, R106–R114 (2016).

99. Villaseñor, R. et al. ChromID identifies the protein interactome at chromatin marks. Nat. Biotechnol. 38, 728–736 (2020).

100. Stieglmeier, M. et al. Nitrososphaera viennensis gen. nov., sp. nov., an aerobic and mesophilic, ammonia-oxidizing archaeon from soil and a member of the archaeal phylum Thaumarchaeota. Int. J. Syst. Evol. Microbiol. 64, 2738–2752 (2014).

101. Tirichine, L. et al. Histone extraction protocol from the two model diatoms Phaeodactylum tricornutum and Thalassiosira pseudonana. Mar. Genomics 13, 21–25 (2014).

102. Shechter, D., Dormann, H. L., Allis, C. D. & Hake, S. B. Extraction, purification and analysis of histones. Nat. Protoc. 2, 1445–57 (2007).

103. Perkins, D. N., Pappin, D. J. C., Creasy, D. M. & Cottrell, J. S. Probability-based protein identification by searching sequence databases using mass spectrometry data. Electrophoresis 20, 3551–3567 (1999).

104. Taus, T. et al. Universal and confident phosphorylation site localization using phosphoRS. J. Proteome Res. 10, 5354–5362 (2011).

105. Vizcaíno, J. A. et al. 2016 update of the PRIDE database and its related tools. Nucleic Acids Res. 44, D447–56 (2016).

106. Hagberg, A. A., Schult, D. A. & Swart, P. J. Exploring Network Structure, Dynamics, and Function using NetworkX. in Proceedings of the 7th Python in Science Conference (eds. Varoquaux, G., Vaught, T. & Millman, J.) 11–15 (2008).

107. Katoh, K. & Standley, D. M. MAFFT Multiple Sequence Alignment Software Version 7: Improvements in Performance and Usability. Mol. Biol. Evol. 30, 772–780 (2013).

108. Nguyen, L.-T., Schmidt, H. A., von Haeseler, A. & Minh, B. Q. IQ-TREE: A Fast and Effective Stochastic Algorithm for Estimating Maximum-Likelihood Phylogenies. Mol. Biol. Evol. 32, 268–274 (2015).

109. Veluchamy, A. et al. An integrative analysis of post-translational histone modifications in the marine diatom Phaeodactylum tricornutum. Genome Biol. 16, 102 (2015).

110. Ren, Q. & Gorovsky, M. a. Histone H2A.Z acetylation modulates an essential charge patch. Mol. Cell 7, 1329–35 (2001).

111. Allis, C. D. et al. hv1 is an evolutionarily conserved H2A variant that is preferentially associated with active genes. J. Biol. Chem. 261, 1941–1948 (1986).

112. Fusauchi, Y. & Iwai, K. Tetrahymena histone H2A. Acetylation in the N-terminal sequence and phosphorylation in the C-terminal sequence. J. Biochem. 95, 147–154 (1984).

113. Xiong, L., Adhvaryu, K. K., Selker, E. U. & Wang, Y. Mapping of Lysine Methylation and Acetylation in Core Histones of Neurospora crassa. Biochemistry 49, 5236–5243 (2010).

114. Zhang, K., Sridhar, V. V., Zhu, J., Kapoor, A. & Zhu, J. K. Distinctive core histone post-translational modification patterns in Arabidopsis thaliana. PLoS One 2, (2007).

115. Johnson, L. et al. Mass spectrometry analysis of Arabidopsis histone H3 reveals distinct combinations of post-translational modifications. Nucleic Acids Res. 32, 6511–6518 (2004).

116. Bergmüller, E., Gehrig, P. M. & Gruissem, W. Characterization of post-translational modifications of histone H2B-variants isolated from Arabidopsis thaliana. J. Proteome Res. 6, 3655–3668 (2007).

117. Beck, H. C. et al. Quantitative Proteomic Analysis of Post-translational Modifications of Human Histones. Mol. Cell. Proteomics 5, 1314–1325 (2006).

118. Goudarzi, A. et al. Dynamic Competing Histone H4 K5K8 Acetylation and Butyrylation Are Hallmarks of Highly Active Gene Promoters. Mol. Cell 62, 169–180 (2016).

119. Hake, S. B. et al. Expression patterns and post-translational modifications associated with mammalian histone H3 variants. J. Biol. Chem. 281, 559–568 (2006).

120. Tan, M. et al. Identification of 67 histone marks and histone lysine crotonylation as a new type of histone modification. Cell 146, 1016–1028 (2011).

121. Moniruzzaman, M., Martinez-Gutierrez, C. A., Weinheimer, A. R. & Aylward, F. O. Dynamic genome evolution and complex virocell metabolism of globally-distributed giant viruses. Nat. Commun. 11, 1–11 (2020).

122. Punta, M. et al. The Pfam protein families database. Nucleic Acids Res. 40, D290–301 (2012).

123. Eddy, S. R. Accelerated profile HMM searches. PLoS Comput. Biol. 7, e1002195 (2011).

124. Buchfink, B., Xie, C. & Huson, D. H. Fast and sensitive protein alignment using DIAMOND. Nat. Methods 12, 59–60 (2015).

125. Enright, A. J., Van Dongen, S. & Ouzounis, C. A. An efficient algorithm for large-scale detection of protein families. Nucleic Acids Res. 30, 1575–1584 (2002).

126. Steenwyk, J. L., Buida, T. J., Li, Y., Shen, X.-X. & Rokas, A. ClipKIT: A multiple sequence alignment trimming software for accurate phylogenomic inference. PLOS Biol. 18, e3001007 (2020).

127. Minh, B. Q., Nguyen, M. A. T. & von Haeseler, A. Ultrafast approximation for phylogenetic bootstrap. Mol. Biol. Evol. 30, 1188–95 (2013).

128. Grau-Bové, X. & Sebé-Pedrós, A. Orthology Clusters from Gene Trees with Possvm. Mol. Biol. Evol. 38, 5204–5208 (2021).

129. Huerta-Cepas, J., Dopazo, H., Dopazo, J. & Gabaldón, T. The human phylome. Genome Biol. 8, R109 (2007).

130. Huerta-Cepas, J., Serra, F. & Bork, P. ETE 3: Reconstruction, Analysis, and Visualization of Phylogenomic Data. Mol. Biol. Evol. 33, 1635–1638 (2016).

131. Csűrös, M. & Miklós, I. A Probabilistic Model for Gene Content Evolution with Duplication, Loss, and Horizontal Transfer BT - Research in Computational Molecular Biology. in (eds. Apostolico, A., Guerra, C., Istrail, S., Pevzner, P. A. & Waterman, M.) 206–220 (Springer Berlin Heidelberg, 2006).

132. Csurös, M. Count: evolutionary analysis of phylogenetic profiles with parsimony and likelihood. Bioinformatics 26, 1910–2 (2010).

133. Gansner, E. R. & North, S. C. An open graph visualization system and its applications to software engineering. Softw. Pract. Exp. 30, 1203–1233 (2000).

134. Fu, L., Niu, B., Zhu, Z., Wu, S. & Li, W. CD-HIT: accelerated for clustering the next-generation sequencing data. Bioinformatics 28, 3150–3152 (2012).

135. Jombart, T., Balloux, F. & Dray, S. adephylo: new tools for investigating the phylogenetic signal in biological traits. Bioinformatics 26, 1907–1909 (2010).

136. Wells, J. N. & Feschotte, C. A Field Guide to Eukaryotic Transposable Elements. Annu. Rev. Genet. 54, annurev-genet-040620-022145 (2020).

137. Storer, J., Hubley, R., Rosen, J., Wheeler, T. J. & Smit, A. F. The Dfam community resource of transposable element families, sequence models, and genome annotations. Mob. DNA 12, 2 (2021).

138. Camacho, C. et al. BLAST+: architecture and applications. BMC Bioinformatics 10, 421 (2009).

139. Li, H. & Durbin, R. Fast and accurate short read alignment with Burrows–Wheeler transform. Bioinformatics 25, 1754–1760 (2009).

140. Quinlan, A. R. & Hall, I. M. BEDTools: a flexible suite of utilities for comparing genomic features. Bioinforma. 26, 841–842 (2010).

141. Anisimova, M., Gil, M., Dufayard, J.-F., Dessimoz, C. & Gascuel, O. Survey of Branch Support Methods Demonstrates Accuracy, Power, and Robustness of Fast Likelihood-based Approximation Schemes. Syst. Biol. 60, 685–699 (2011).

142. Bodenhofer, U., Bonatesta, E., Horejš-Kainrath, C. & Hochreiter, S. msa: an R package for multiple sequence alignment. Bioinformatics 31, 3997–3999 (2015).

