## Supplementary figures and images for "A phylogenetic and proteomic reconstruction of eukaryotic chromatin evolution"

### species_tree.newick.pdf

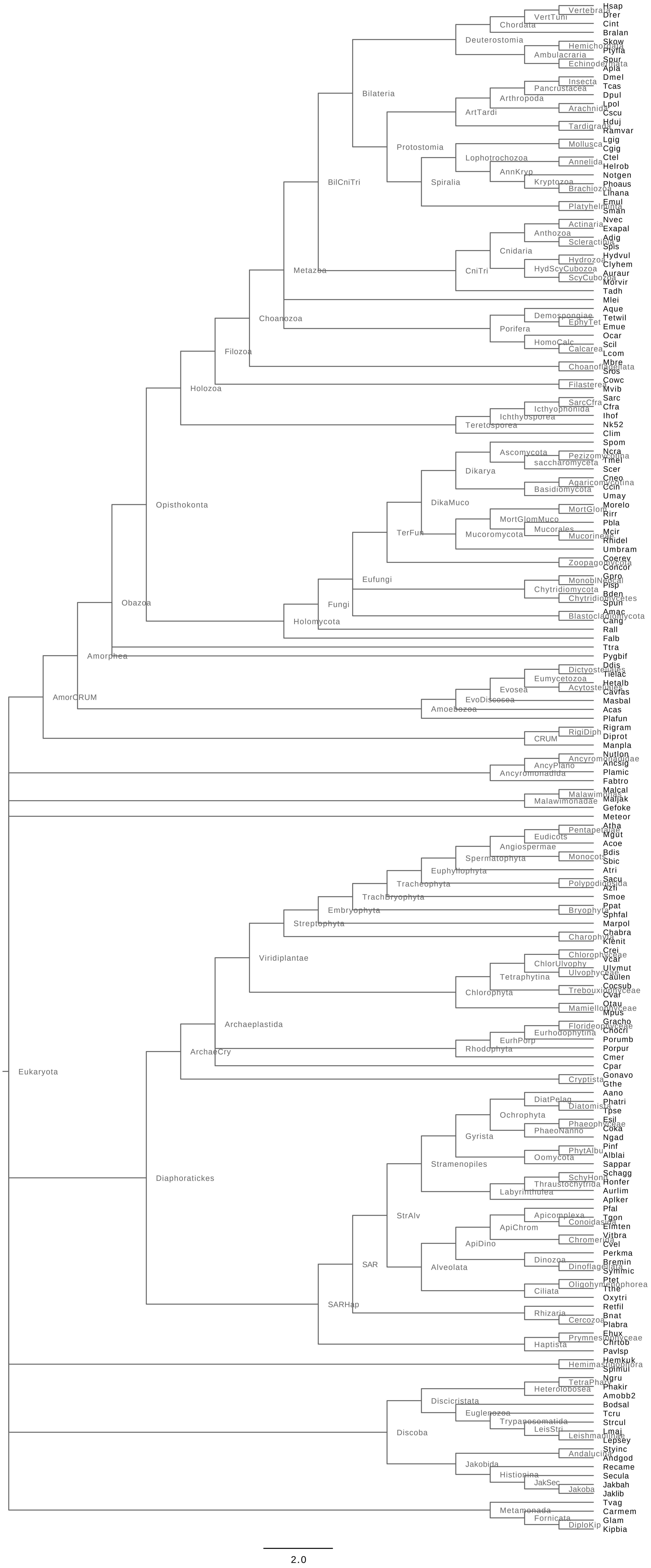

### Supplementary Fig. 1

##### Supplementary Figure S1

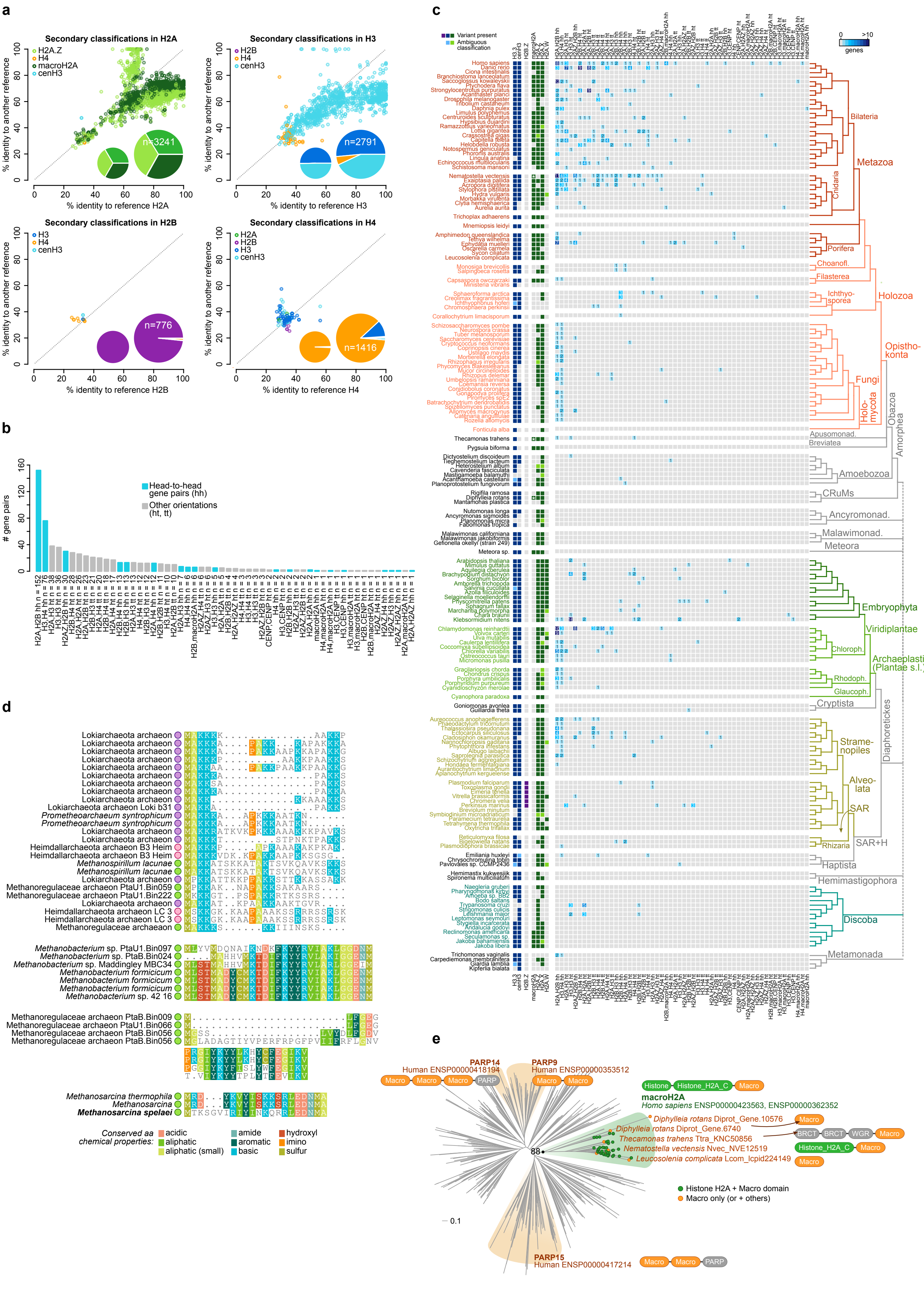

### Supplementary Fig. 2

#### Supplementary Figure S2

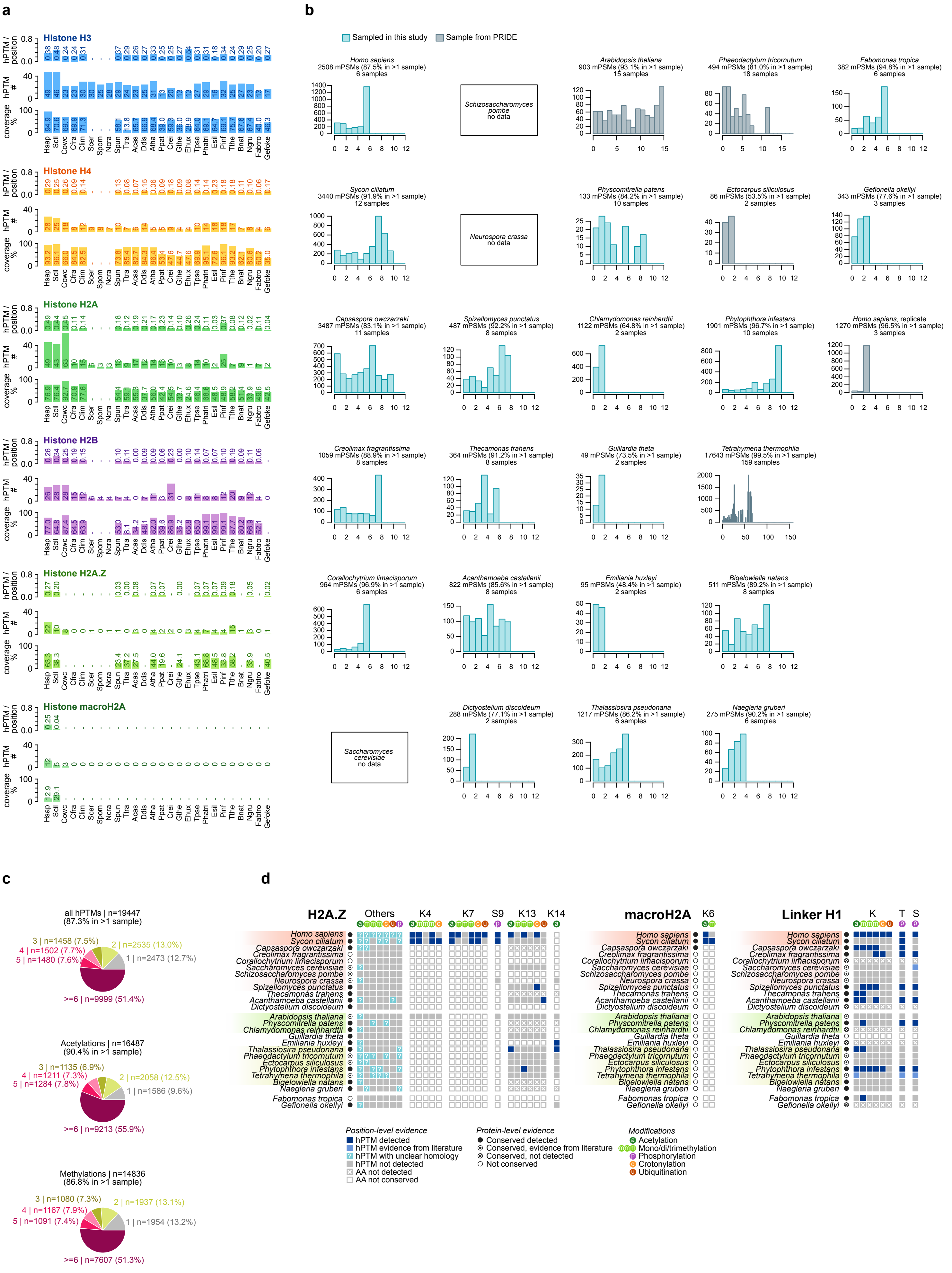

### Supplementary Fig. 3

**Supplementary Figure S3**

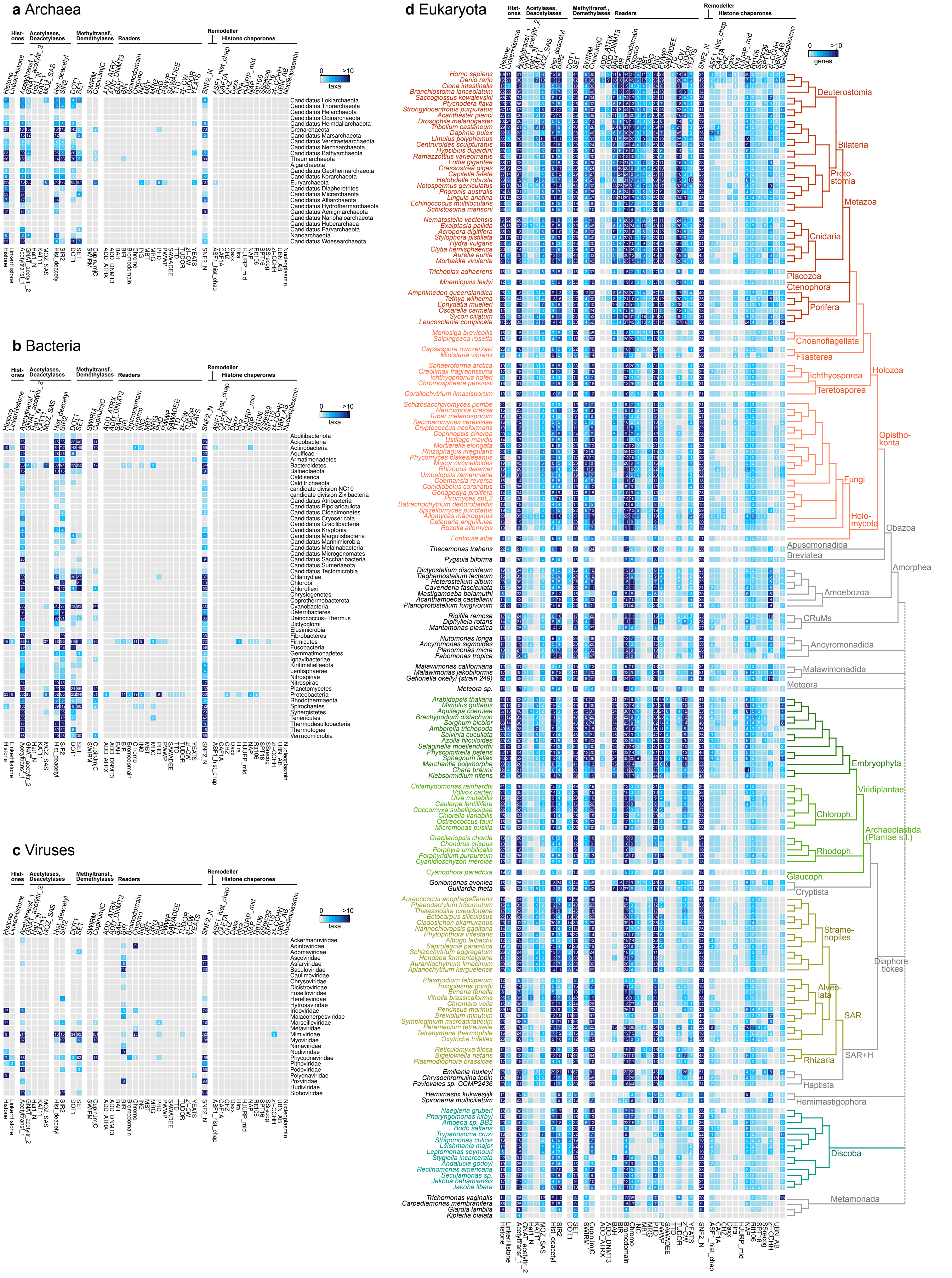

### Supplementary Fig. 4

##### Supplementary Figure S4

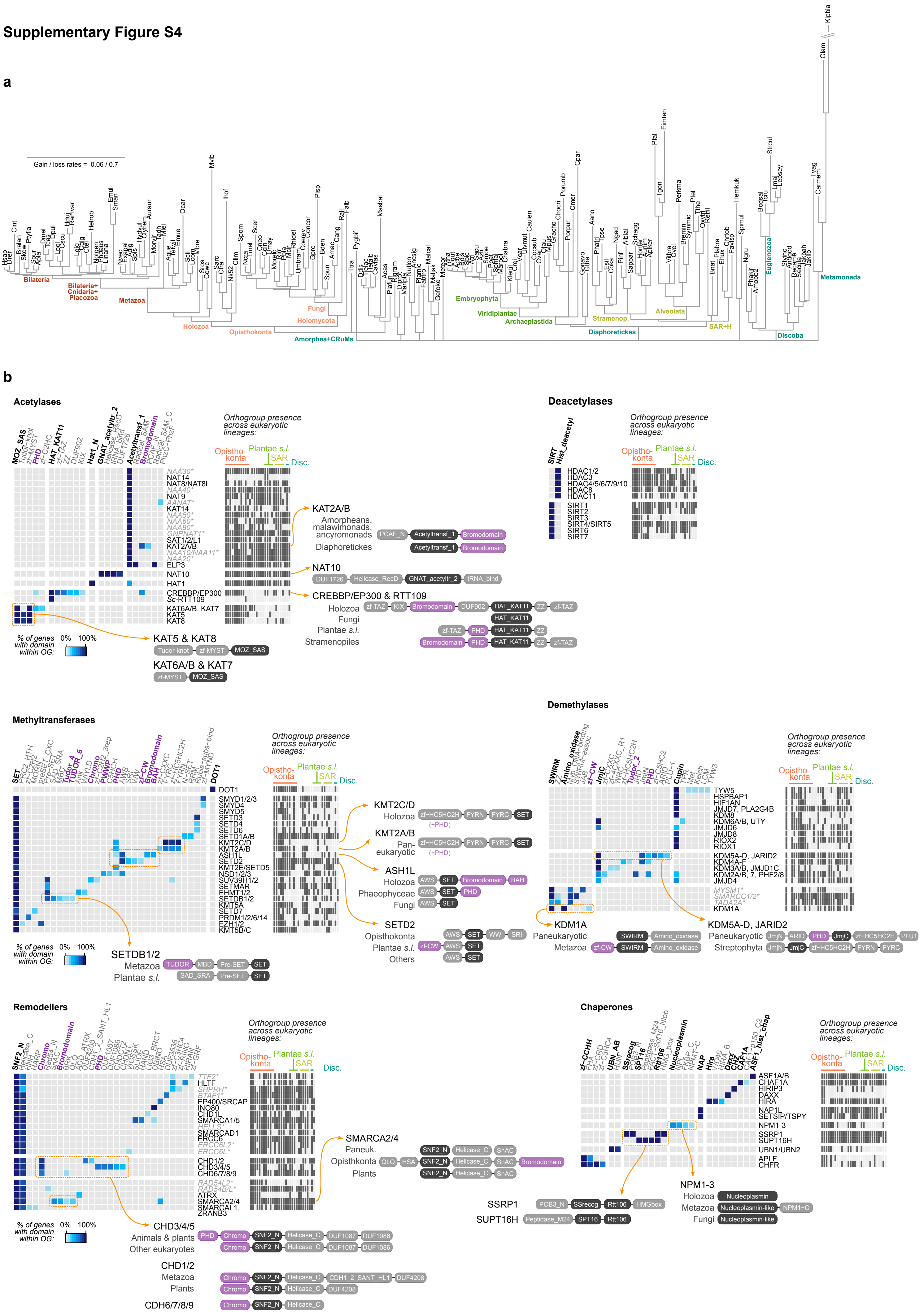

### Supplementary Fig. 5

#### Supplementary Figure S5

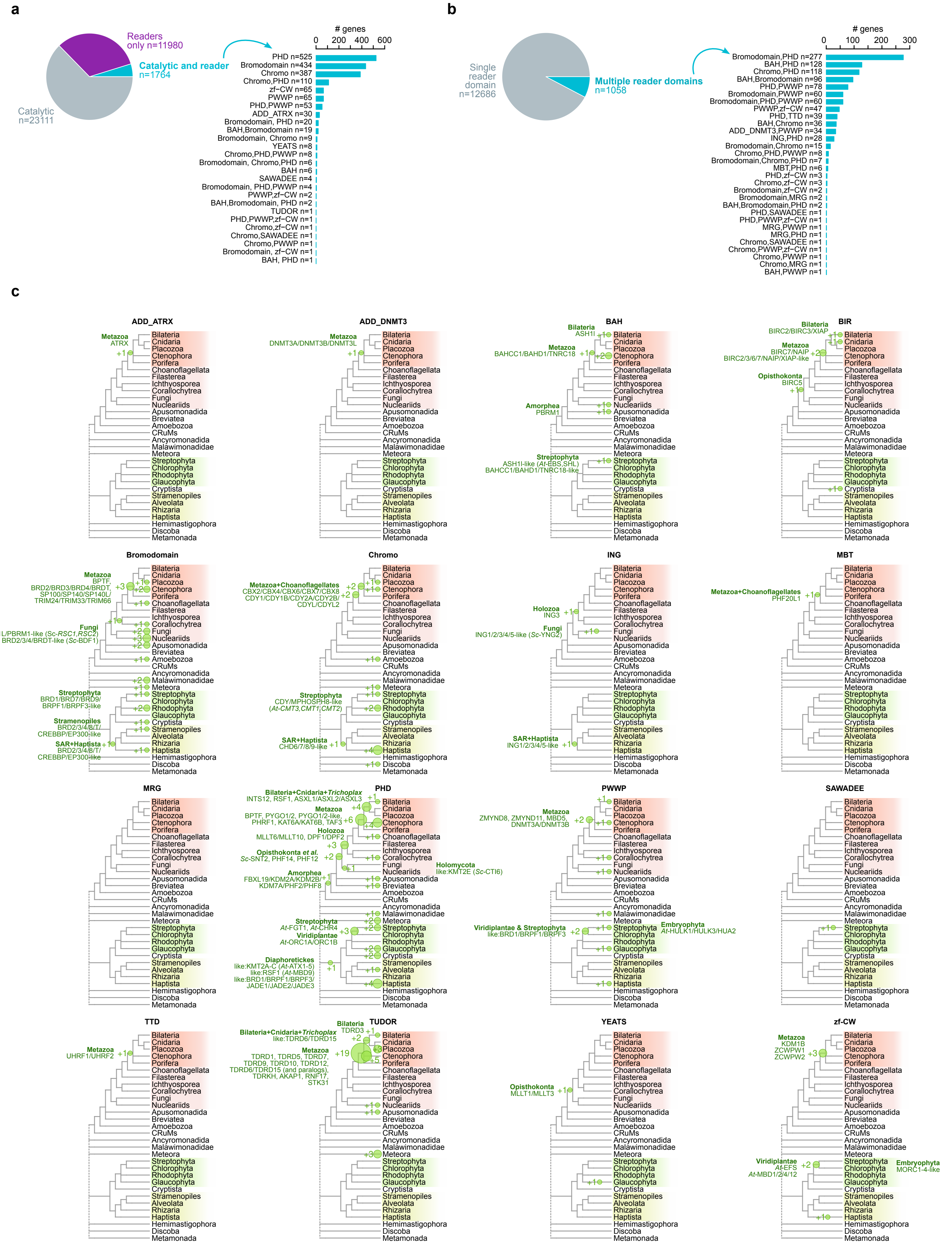

### Supplementary Fig. 6

##### Supplementary Figure S6

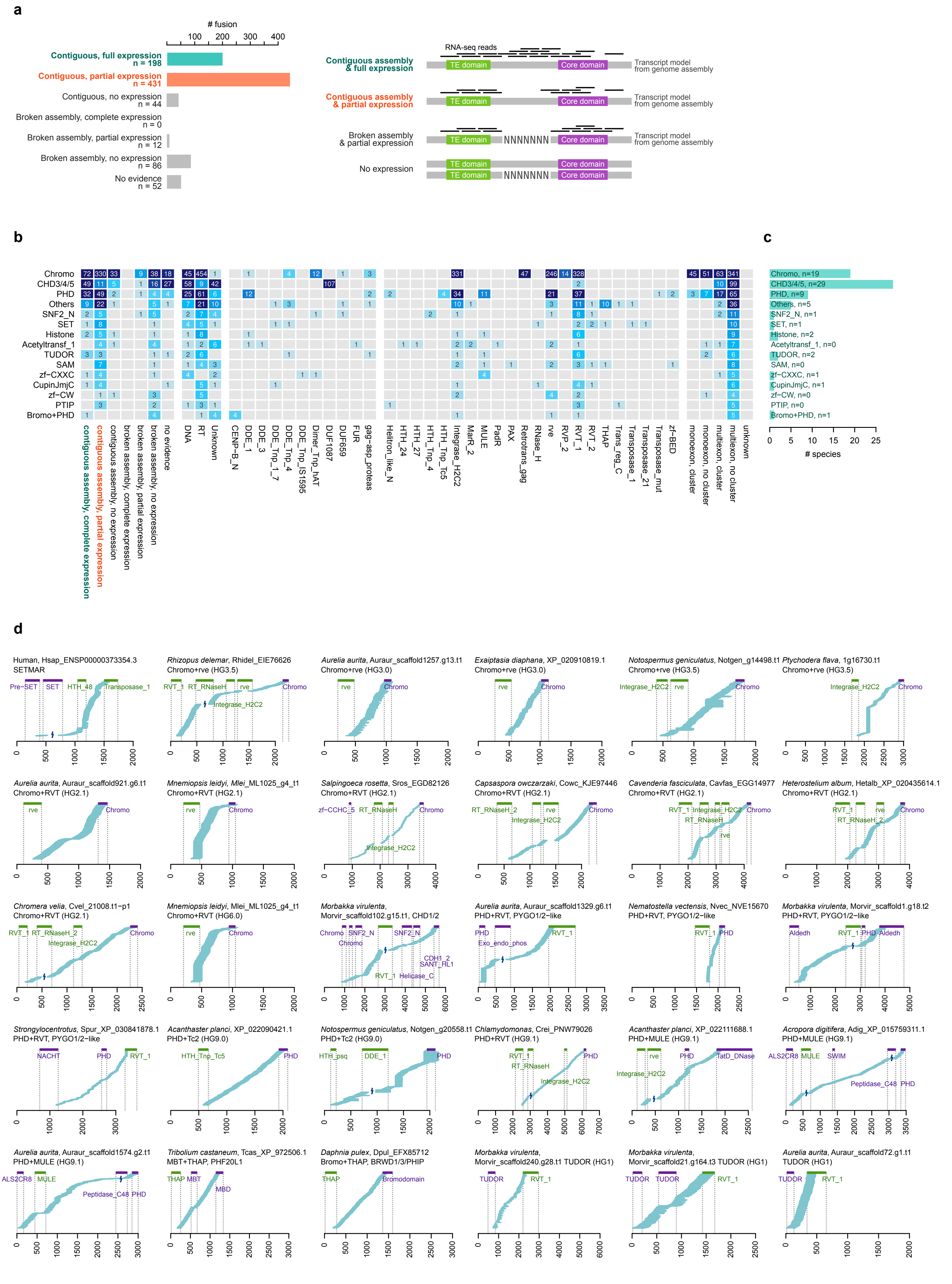

### Supplementary Fig. 7

##### Supplementary Figure S7

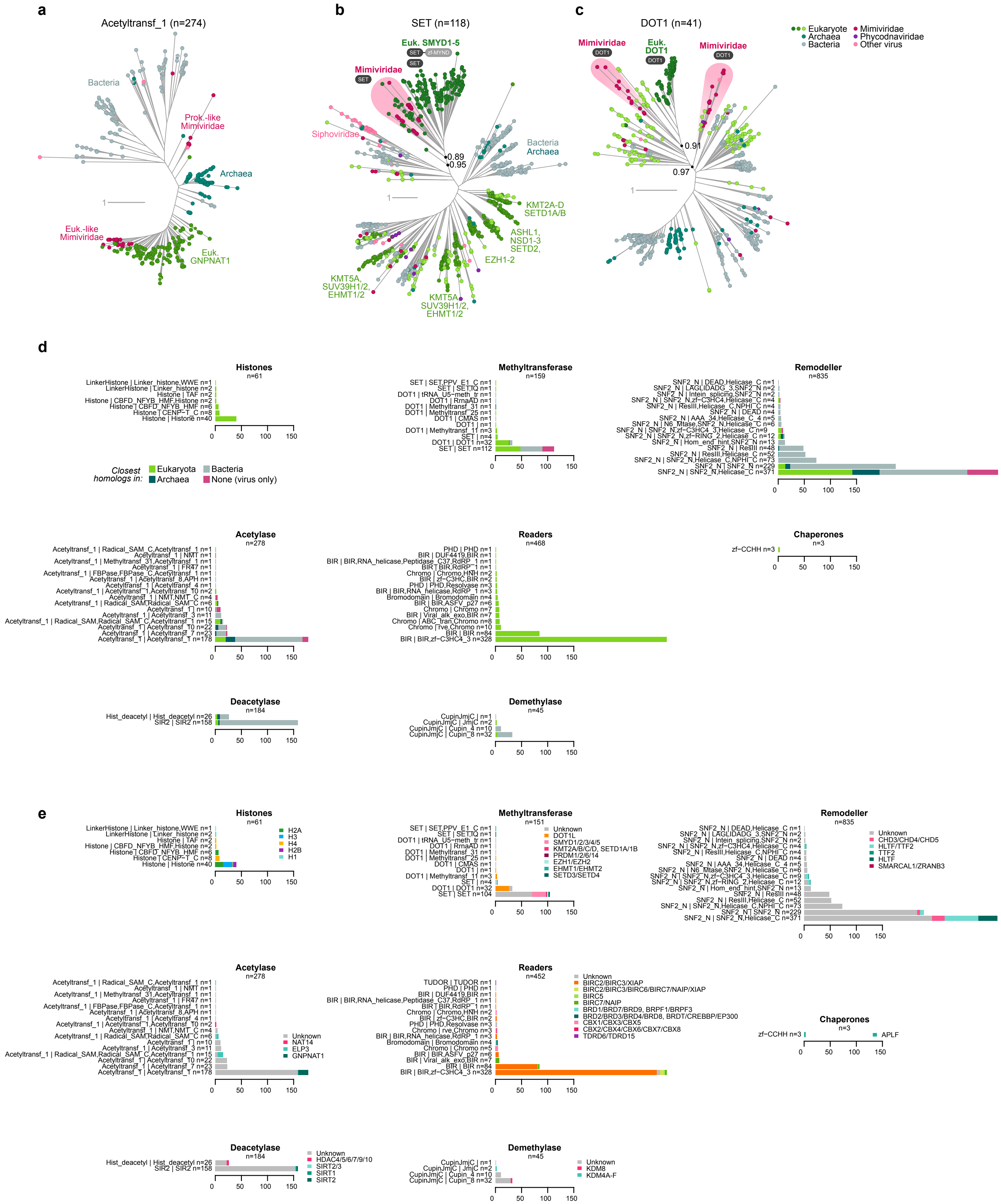
